# Systemic LSD1 inhibition prevents aberrant remodeling of metabolism in obesity

**DOI:** 10.1101/2021.11.25.469928

**Authors:** Bastian Ramms, Dennis P. Pollow, Han Zhu, Chelsea Nora, Austin R. Harrington, Ibrahim Omar, Philip L.S.M. Gordts, Matthew Wortham, Maike Sander

## Abstract

The transition from lean to obese states involves systemic metabolic remodeling that impacts insulin sensitivity, lipid partitioning, inflammation, and glycemic control. Here, we have taken a pharmacological approach to test the role of a nutrient-regulated chromatin modifier, lysine-specific demethylase (LSD1), in obesity-associated metabolic reprogramming. We show that systemic administration of an LSD1 inhibitor (GSK-LSD1) reduces food intake and body weight, ameliorates non-alcoholic fatty liver disease (NAFLD), and improves insulin sensitivity and glycemic control in mouse models of obesity. GSK-LSD1 has little effect on systemic metabolism of lean mice, suggesting LSD1 has a context-dependent role in promoting maladaptive changes in obesity. Analysis of insulin target tissues identified white adipose tissue as the major site of insulin sensitization by GSK-LSD1, where it reduces adipocyte inflammation and lipolysis. We demonstrate that GSK-LSD1 reverses NAFLD in a non-hepatocyte-autonomous manner, suggesting an indirect mechanism via inhibition of adipocyte lipolysis and subsequent effects on lipid partitioning. Pair-feeding experiments further revealed that effects of GSK-LSD1 on hyperglycemia and NAFLD are not a consequence of reduced food intake and weight loss. These findings suggest that targeting LSD1 could be a strategy for treatment of obesity and its associated complications including type 2 diabetes and NAFLD.

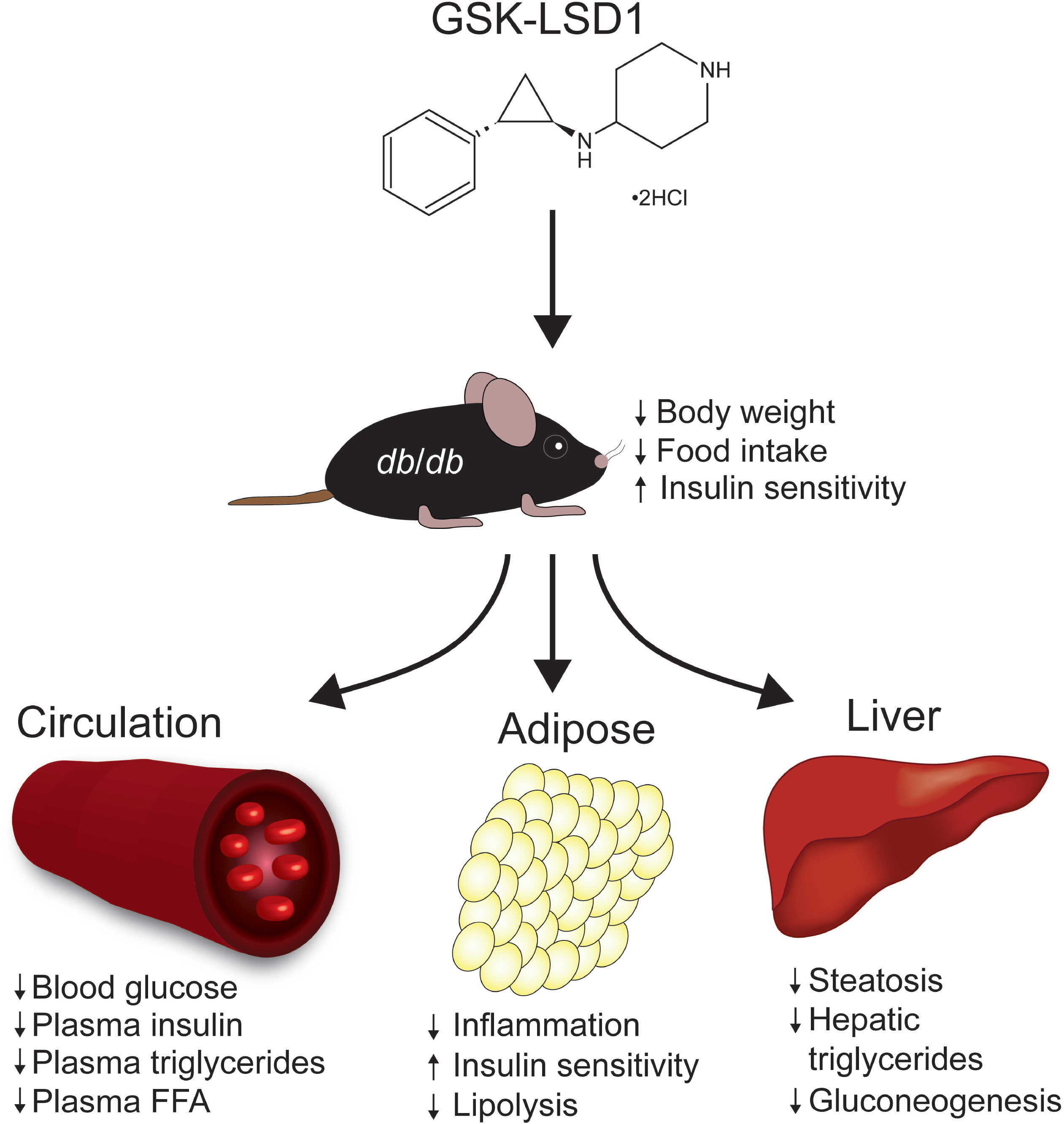

## Introduction

The root cause of obesity is an imbalance between energy intake and expenditure. The resulting body weight gain is associated with remodeling of systemic metabolism. The metabolic remodeling gives rise to interrelated defects including insulin resistance, ectopic lipid deposition, and chronic, low-grade inflammation (1, 2). These defects predispose people with obesity to more serious chronic diseases including cardiovascular disease, type 2 diabetes (T2D), and nonalcoholic fatty liver disease (NAFLD), which together comprise much of the public health burden of obesity.

Obesity causes insulin resistance in white adipose tissue (WAT) (3, 4), thereby increasing lipolysis and the trafficking of free fatty acids (FFA) from WAT to the liver (5, 6). This oversupply of FFA to the liver contributes to accumulation of triglycerides that promote hepatic steatosis, the defining characteristic of NAFLD (6–8). Systemic insulin resistance caused by obesity increases pancreatic β-cell workload and thereby also predisposes to T2D, which manifests when the β-cell adaptive response fails to produce adequate insulin to meet demand (6, 9). The hyperinsulinemia associated with successful β-cell adaptation promotes lipogenesis in the liver and exacerbates hepatic steatosis, underscoring the systemic nature of metabolic defects in obesity (10, 11). Despite the urgent global need to ameliorate complications associated with obesity, there are few therapeutic options to treat obesity itself (2). As obesity is a complex multi-organ disease, therapies that address its root cause or simultaneously target the secondary metabolic defects may be necessary to mitigate the associated complications. Unfortunately, current therapeutic efforts for obesity are limited by an incomplete understanding of how the nutrient environment is interpreted by various metabolically relevant tissues to evoke adaptive or maladaptive changes in obesity (12). Overall, there is an unmet need to understand context-dependent regulation of metabolism in the obese compared to the lean state.

Adaptation of systemic metabolism to changing nutrient states involves several layers of regulation that affect the intake, storage, and expenditure of energy. These processes are controlled at the whole-organism level through the endocrine and central nervous systems (13, 14). It is currently unclear whether these systemic controls are augmented by unified tissue-intrinsic mechanisms. One such potential mechanism is the transcriptional regulation of metabolism, which is mediated by transcription factors and coregulators whose functions are coupled to nutrient state (15–19). For example, intermediary metabolites have been shown to regulate the enzymatic activities of several coregulators that modify the epigenome (20, 21). Changes in cellular metabolism lead to altered abundance of substrates for these coregulators, thereby altering their enzymatic activities to change epigenomic states and gene expression (22). Posttranslational modifications of coregulators in response to hormonal or nutrient signals provide an additional layer of regulation that links coregulator function to nutrient state (15, 17). Genetic inactivation of transcriptional coregulators has revealed pervasive roles for these enzymes in governing nutrient responses of metabolically relevant tissues (15-18, 23-27). For instance, SIN3A is an insulin-regulated corepressor of the glucokinase gene that restrains hepatic lipogenesis in the fasted state. In the absence of SIN3A, the lipogenic effect of insulin is dampened (18). Overall, nutrient regulation confers context-specific functions to transcriptional coregulators, and therefore these enzymes could underlie context-dependent regulatory mechanisms specific to lean or obese states. Given recent drug development efforts to target transcriptional coregulators, understanding how the compendium of these enzymes impacts metabolism could lead to new therapies for metabolic disease.

We and others have identified key roles for lysine-specific histone demethylase 1A (LSD1/KDM1A) in context-dependent regulation of metabolism. The demethylase activity of LSD1 requires flavin adenine dinucleotide (FAD) as a cofactor, linking cellular metabolism to the nutrient environment (26). Given its nutrient responsiveness and broad expression pattern (28), we reasoned that LSD1 would be well-positioned to mediate systemic changes to metabolism. LSD1 is part of transcriptional complexes in the nucleus that regulate nutrient responses in various cell types including hepatocytes, adipocytes, and pancreatic β-cells (21, 27–32). LSD1 restrains the insulin secretory response by pancreatic β-cells in the fed state (27), whereas in the liver, LSD1 promotes insulin-stimulated lipogenesis in response to feeding (29–31). Furthermore, beiging of WAT during cold exposure requires LSD1 to activate thermogenesis (32, 33). While these studies indicate that LSD1 regulates context-specific functions of several metabolically relevant tissues, it is unknown whether LSD1 is involved in the remodeling of systemic metabolism in response to changing nutrient states.

In the current study, we employed a pharmacological approach to test the role of LSD1 in metabolic remodeling associated with obesity. We show that systemic LSD1 inhibition reduces hyperphagia and weight gain, improves insulin sensitivity, and prevents hyperglycemia in obesity models, while having no discernable metabolic effects in lean mice. Mechanistically, we found that LSD1 inhibition reduces adipose tissue inflammation and ameliorates NAFLD. In intervention studies, we demonstrate that LSD1 inhibition can reverse Western diet-induced weight gain and hyperinsulinemia as well as enhance insulin signaling. Together, our data suggest that LSD1 could be a therapeutic target in metabolic disease that simultaneously addresses the root cause of obesity and its associated complications.

## Results

### LSD1 inhibition prevents hyperglycemia in db/db mice

To investigate whether LSD1 contributes to metabolic dysfunction in obesity, we studied the effect of systemic LSD1 inhibition in *db*/*db* mice. Four-week-old *db*/*db* mice and lean *db*/+ control mice were injected daily with GSK-LSD1, an LSD1 inhibitor, or vehicle (veh) and metabolic parameters were monitored longitudinally for six weeks (**Figure 1A**). As expected (34), veh-treated *db*/*db* mice rapidly gained weight and became hyperglycemic around six weeks of age (**Figure 1B**). GSK-LSD1 reduced weight gain of *db*/*db* mice (**Figure 1B**). Remarkably, blood glucose levels of GSK-LSD1-treated *db*/*db* mice were comparable to levels in lean control mice (**Figure 1C**). GSK-LSD1-treated *db*/*db* mice also exhibited improved glucose tolerance (**Figure 1D**). Notably, GSK-LSD1 had no effect on body weight or blood glucose levels in lean *db*/+ mice (**Supplemental Figure 1A** and **B**), suggesting that LSD1 regulates maladaptive changes to metabolism in obesity. Together, these findings show that systemic LSD1 inhibition prevents development of diabetes in *db*/*db* mice, a model of obesity and T2D.

**Figure 1.**
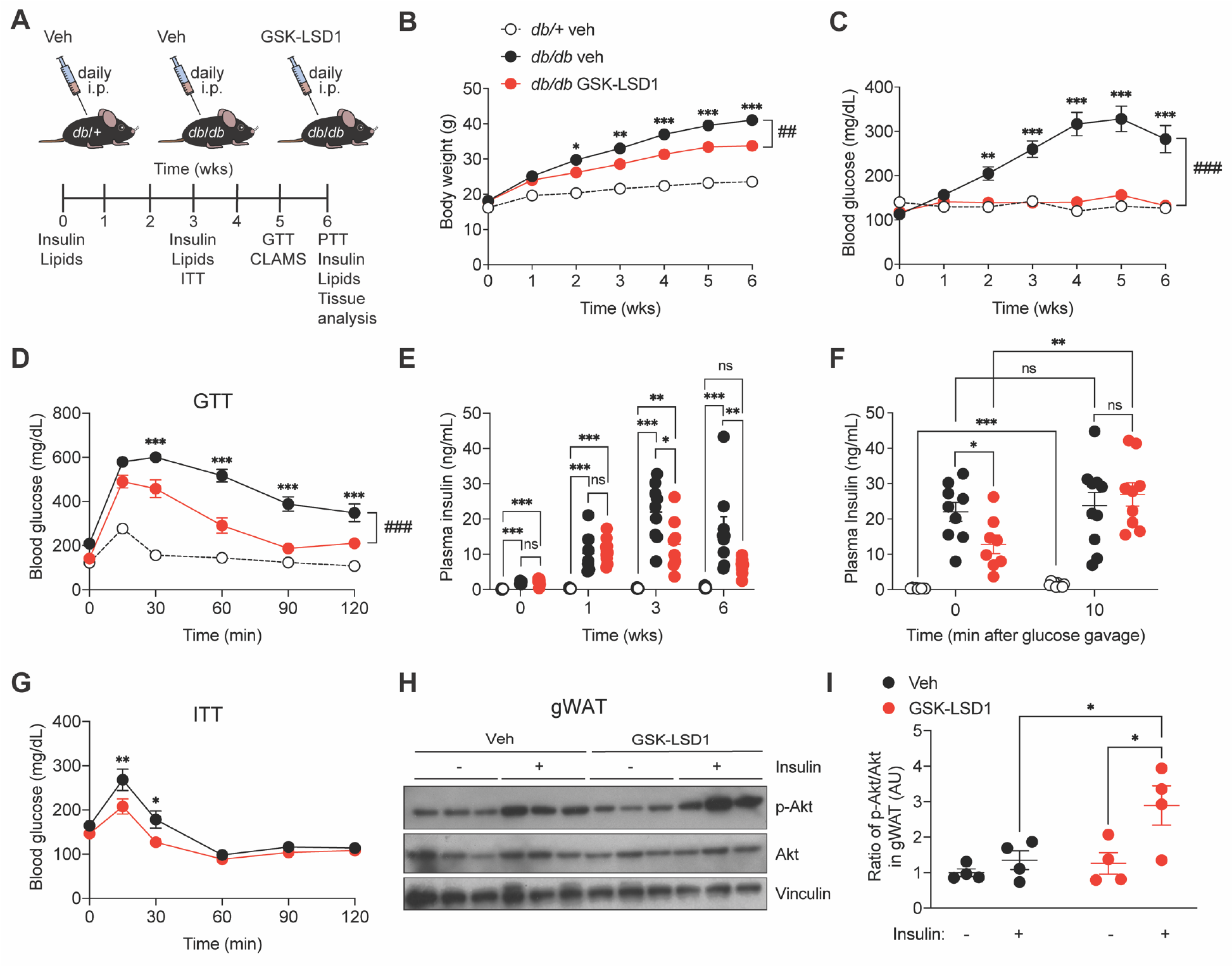
Systemic LSD1 inhibition prevents the development of hyperglycemia and improves insulin sensitivity in *db*/*db* mice. (**A**) 4-week-old *db*/*db* mice received daily intraperitoneal (i.p.) injections of GSK-LSD1 or vehicle (veh) for 6 weeks. As a control, lean *db*/+ mice were injected with veh. Metabolic measurements were conducted at the indicated time points. (**B**) Body weight and (**C**) blood glucose levels measured weekly (n = 10 mice/group). (**D**) Blood glucose levels at indicated time points after a glucose bolus via oral gavage (n = 10 mice/group). Glucose tolerance test, GTT. (**E**) Fasting plasma insulin levels at baseline and after 1, 3, and 6 weeks of GSK-LSD1 or veh treatment (n = 6-10 mice/group). (**F**) Plasma insulin levels before and 10 min after glucose gavage (n = 8-10 mice/group). (**G**) Blood glucose levels at indicated time points after intraperitoneal insulin (2.0 U/kg body weight) injection (n = 10 mice/group). Insulin tolerance test, ITT. (**H**) Immunoblot analysis of pAkt^Ser473^, Akt, and vinculin in gWAT. GSK-LSD1-or veh-treated *db*/*db* mice injected with insulin or saline. (**I**) Quantification of pAkt^Ser473^ to Akt ratio as fold change compared to veh-treated mice without insulin injection (n = 4 mice/group). Data are presented as mean ± SEM. Statistical differences were calculated using a two-way ANOVA with Tukey post hoc analysis (in B-D, F, G, I). A one-way ANOVA with Tukey post hoc analysis was performed to analyze statistical differences between three or more groups (E). Unless otherwise indicated, significance is shown between GSK-LSD1- and veh-treated mice. \**p*<0.05, \*\**p*<0.01, \*\*\**p*<0.001, ^##^*p*<0.01, ^###^*p*<0.001. AU, arbitrary unit; ns, not significant.

To identify mechanisms leading to improved glycemic control after GSK-LSD1 treatment, we measured plasma insulin levels and performed insulin tolerance tests in veh- and GSK-LSD1-treated *db*/*db* mice. Fasting plasma insulin levels were significantly reduced after three and six weeks of GSK-LSD1 treatment (**Figure 1E**). Moreover, GSK-LSD1-treated mice showed a two-fold increase in plasma insulin levels after glucose stimulation compared to baseline, whereas glucose stimulation did not significantly increase insulin levels in the veh group (**Figure 1F**), indicating improved coupling of insulin secretion with blood glucose following LSD1 inhibition. Insulin tolerance tests revealed increased insulin sensitivity in the GSK-LSD1 treatment group compared to veh-treated *db*/*db* mice (**Figure 1G**).

Insulin resistance in peripheral insulin target tissues is a major characteristic of obesity (6, 35). We therefore assessed signal transduction downstream of the insulin receptor by measuring p-Akt^Ser473^ in WAT, liver, and skeletal muscle following insulin injection of *db*/*db* mice. Insulin-stimulated p-Akt^Ser473^ was higher in gonadial white adipose tissue (gWAT) of the GSK-LSD1 group compared to the veh group (**Figure 1H** and **I**). By contrast, LSD1 inhibition had little effect on p-Akt^Ser473^ in skeletal muscle and liver (**Supplemental Figure 1C-F**). These results suggest that systemic LSD1 inhibition improves glucose homeostasis of *db*/*db* mice in part through enhanced insulin sensitivity of adipose tissue.

### LSD1 inhibition reduces adipose inflammation and lipolysis

To examine effects of GSK-LSD1 on adipose tissue, we first measured adipose tissue weight and adipocyte size in gWAT, subcutaneous white adipose tissue (sWAT), and brown adipose tissue (BAT). The tissue weights of gWAT and sWAT were significant reduced in the GSK-LSD1-treated group, indicating a loss of fat mass following LSD1 inhibition (**Supplemental Figure 2A-C**). The fat mass loss was not a result of reduced adipocyte size (**Figure 2A** and **Supplemental Figure 2D-H**).

**Figure 2.**
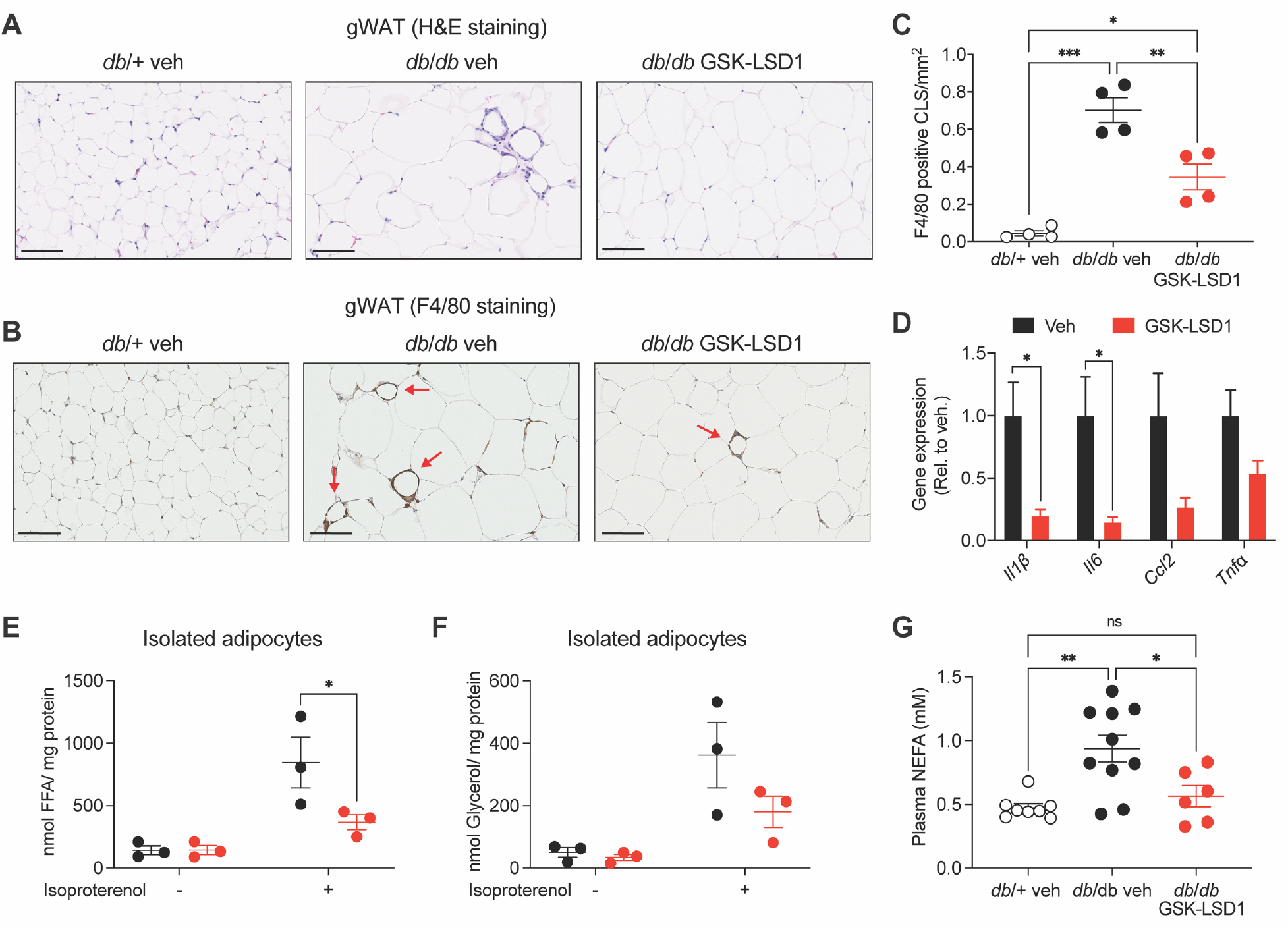
LSD1 inhibition reduces adipose tissue inflammation and lipolysis in *db*/*db* mice. (**A**) Representative images of gWAT sections stained with hematoxylin and eosin (H&E) and (**B**) detection of crown-like structures (CLS) using an F4/80 antibody in gWAT after 6 weeks of daily GSK-LSD1 or vehicle (veh) administration to *db*/*db* mice. As a control, lean *db*/+ mice were injected with veh. Red arrows highlight CLS. Scale bars = 100 µm. (**C**) Quantification of F4/80^+^ CLS in gWAT relative to tissue size (n = 4 mice/group). (**D**) qPCR analysis of inflammatory genes in gWAT. Transcript levels in GSK-LSD1-treated relative to veh-treated *db*/*db* mice (n = 6-11 mice/group). (**E**,**F**) Lipolysis in differentiated adipocytes isolated from *db*/*db* mice after preincubation with GSK-LSD1 or veh and stimulation with isoproterenol. (**E**) Free fatty acid (FFA) release and (**F**) glycerol release (n = 3 mice). (**G**) Plasma non-esterified fatty acid (NEFA) levels (n = 6-10 mice/group). Data are presented as mean ± SEM. A one-way ANOVA with Tukey post hoc analysis was performed to analyze statistical differences between three or more groups (C, G) or multiple unpaired t-tests to determine differences between two groups (D, E-F). \**p*<0.05, \*\**p*<0.01, \*\*\**p*<0.001, ^#^*p*<0.05, ns, not significant.

A key mechanism leading to impaired insulin sensitivity in WAT is local inflammation (1, 3). WAT inflammation is characterized by infiltration of macrophages, which form crown-like structures (CLS) around dead adipocytes (4, 36) and produce chemokines and cytokines that promote adipose tissue insulin resistance (3, 4). Histological evaluation revealed abundant CLS in gWAT from veh-treated *db*/*db* mice more so than in sWAT and BAT (**Figure 2A** and **Supplemental Figure 2D** and **E**). CLS were rare in gWAT from GSK-LSD1-treated *db*/*db* mice (**Figure 2A**), suggesting that GSK-LSD1 reduces macrophage accumulation in gWAT. Quantification of macrophages based on staining for the macrophage-specific epitope F4/80 confirmed reduced macrophage numbers in gWAT after GSK-LSD1 treatment (**Figure 2B** and **C**). In line with this result, gene expression studies revealed downregulation of inflammatory genes, such as *interleukin 1β* (*Il1β*) and *interleukin 6* (*Il6)* in gWAT (**Figure 2D**). These observations indicate that LSD1 inhibition decreases adipose inflammation in obesity. Considering the association between inflammation and adipose tissue insulin resistance (3), the reduction of adipose inflammation by GSK-LSD1 could contribute to the improvement in adipose insulin sensitivity (**Figure 1H** and **I**). To determine whether the LSD1 inhibitor has direct effects on insulin sensitivity at the level of adipocytes, we measured insulin-stimulated p-Akt^Ser473^ following GSK-LSD1 treatment in differentiated adipocytes of *db*/*db* mice (**Supplemental Figure 2I**). In contrast to our observations in vivo, GSK-LSD1 did not alter insulin signaling in vitro in adipocytes (**Supplemental Figure 2J** and **K**). These data suggest that LSD1 inhibition improves adipose tissue insulin sensitivity indirectly.

Lipolysis is tightly regulated by circulating insulin. WAT insulin resistance is associated with augmented hydrolysis of triglycerides, resulting in increased levels of circulating non-esterified fatty acids (NEFA) (35, 37). To test whether GSK-LSD1-mediated improvements in insulin signaling reduce lipolysis, we pre-incubated differentiated adipocytes from *db*/*db* mice with GSK-LSD1 or veh and then treated with isoproterenol, which induces lipolysis by activating β-adrenergic signaling. GSK-LSD1 reduced isoproterenol-stimulated FFA release concomitant with a trend towards reduced glycerol release (**Figure 2E** and **F**). Moreover, GSK-LSD1 reduced plasma NEFA levels in *db*/*db* mice, suggesting that systemic LSD1 inhibition reduces WAT lipolysis in vivo (**Figure 2G**). Together, these observations indicate that LSD1 inhibition corrects several WAT defects associated with obesity including insulin resistance, inflammation, and augmented lipolysis.

### LSD1 inhibition ameliorates liver steatosis in db/db mice

Obesity and insulin resistance are closely associated with NAFLD (6–8). Having observed that GSK-LSD1 prevents weight gain, increases insulin sensitivity, and reduces FFA release in *db/db* mice, we asked whether GSK-LSD1 ameliorates obesity-associated changes in the liver. Livers from veh-treated *db*/*db* mice showed clear morphological and histological signs of steatosis (**Figure 3A** and **B**). In contrast, livers from *db*/*db* mice treated with GSK-LSD1 were grossly and histologically normal, resembling livers from lean *db*/+ mice. In line with these findings, GSK-

**Figure 3.**
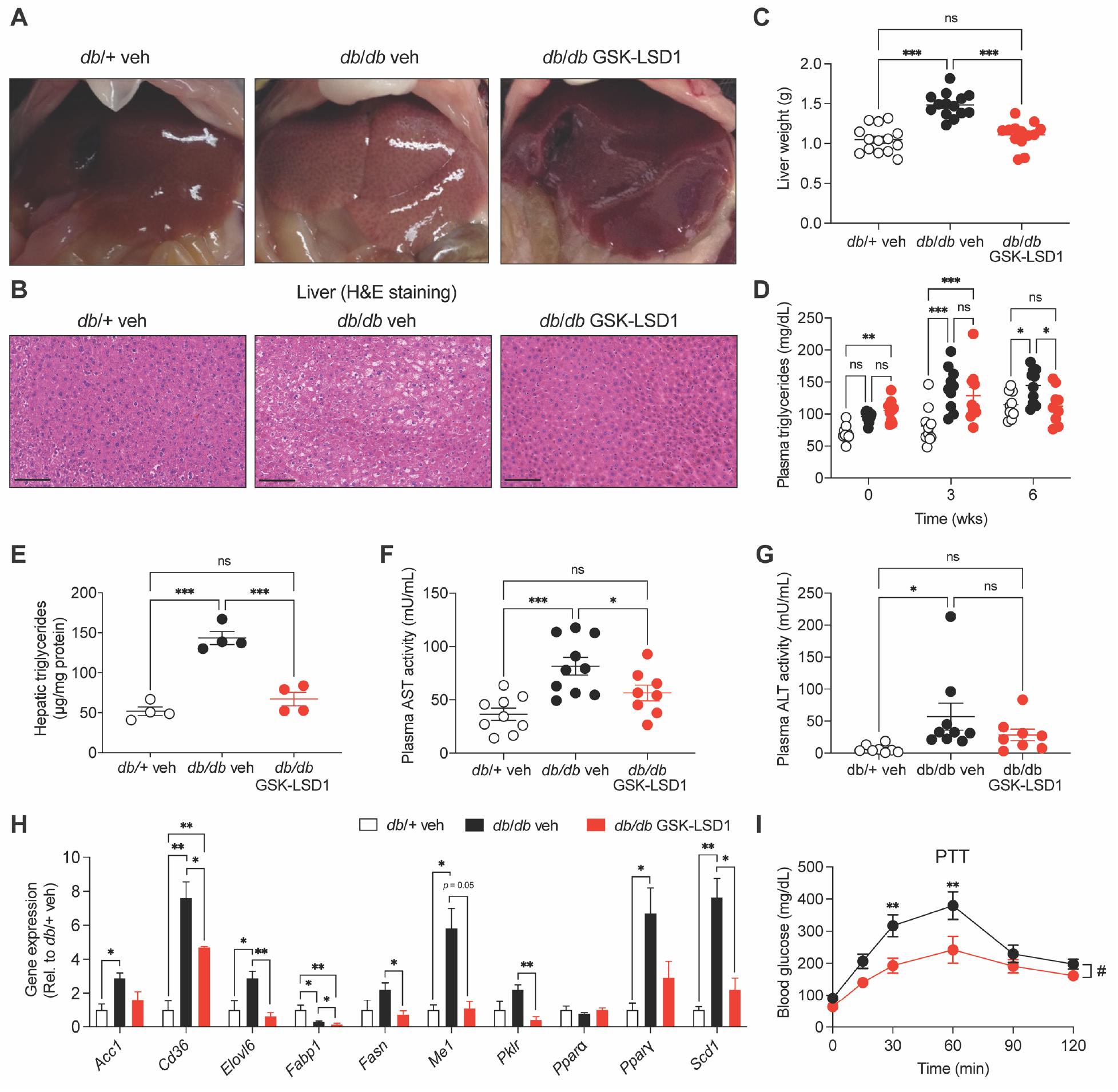
LSD1 inhibition protects against liver steatosis in *db*/*db* mice. (**A**) Representative images of livers in *db*/*db* mice treated daily with GSK-LSD1 or vehicle (veh) for 6 weeks. As a control, lean *db*/+ mice were injected with veh (n = 13-14 mice/group). (**B**) Hematoxylin and eosin (H&E) stain of liver sections. Scale bars = 100 μm. (**C**) Liver weight (n = 13-14 mice/group). (**D**) Fasting plasma triglyceride levels at indicated time points (n = 9-10 mice/group). (**E**) Hepatic triglyceride levels (n = 4 mice/group). (**F**) AST and (**G**) ALT activity in plasma of GSK-LSD1- or veh-treated mice for 6 weeks (n = 8-10 mice/group). (**H**) qPCR analysis of genes associated with hepatic lipid metabolism in liver. Transcript levels relative to *db*/+ mice. (n = 3-4 mice/group). (**I**) Blood glucose levels at indicated time points after intraperitoneal pyruvate injection (n = 10 mice/group). Pyruvate tolerance test, PTT. Data are presented as mean ± SEM. Statistical differences were calculated using a one-way ANOVA (C, E-H) or two-way ANOVA (D, I) with Tukey post hoc analysis. \**p*<0.05, \*\**p*<0.01, \*\*\**p*<0.001, ns, not significant.

LSD1 significantly reduced liver weight in *db*/*db* mice (**Figure 3C**). Steatosis results from a combination of increased lipid delivery from the circulation and increased lipogenesis by hepatocytes, leading to excess intrahepatic accumulation of lipids (7, 8). LSD1 inhibition in *db*/*db* mice decreased both circulating and hepatic triglycerides to levels comparable to those of lean *db*/+ mice (**Figure 3D** and **E**). Of note, plasma and hepatic cholesterol levels as well as circulating levels of the ketone body β-hydroxybutyrate were unaffected GSK-LSD1 administration (**Supplemental Figure 3A-C**). To determine whether GSK-LSD1 improves clinical biomarkers of NAFLD, we measured plasma levels of aspartate aminotransferase (AST) and alanine aminotransferase (ALT). Treatment of *db*/*db* mice with GSK-LSD1 led to a significant reduction in AST activity and a slight, albeit not significant decrease in ALT activity (**Figure 3F** and **G**). These results indicate that systemic LSD1 inhibition prevents liver pathology caused by obesity. In line with the GSK-LSD1-mediated reduction of steatosis, lipogenic genes such as elongation of very long chain fatty acids protein 6 (*Elovl6*), fatty acid synthase (*Fasn*), and stearyl-CoA desasturase (*Scd1*) were downregulated by GSK-LSD1 treatment (**Figure 3H**). LSD1 inhibition further reduced expression of pyruvate kinase isoenzyme R/L (*Pklr*), which catalyzes the last step in the glycolytic pathway and is required for *de novo* lipogenesis from glucose. Moreover, lower expression of fatty acid binding protein 1 (*Fabp1*) and cluster of differentiation 36 (*Cd36*) indicates a possible effect of GSK-LSD1 on liver fatty acid uptake in *db/db* mice. Taken together, these observations suggest that systemic LSD1 inhibition impacts both lipid partitioning and hepatocyte metabolic pathways to ameliorate liver steatosis in obesity.

As the liver is the main site of *de novo* glucose production, we also determined whether GSK-LSD1 reduces hyperglycemia in part through effects on hepatic gluconeogenesis. *db/db* mice that received GSK-LSD1 exhibited lower glucose excursions following pyruvate challenge compared to veh-treated mice (**Figure 3I**), suggesting that systemic LSD1 inhibition improves glucose homeostasis in part through reduced hepatic glucose production.

### Hepatocyte-specific *Lsd1* deletion does not improve glycemia or liver steatosis in *db/db* mice

We reasoned that a reduction in steatosis following GSK-LSD1 treatment could result from altered lipid partitioning from storage tissues or from cell autonomous effects of LSD1 in hepatocytes. While reduced circulating NEFA and triglycerides following LSD1 inhibition supports a potential effect upon lipid partitioning (**Figure 2G** and **3D**), LSD1 is expressed in hepatocytes (**Supplemental Figure 4A**) and could have hepatocyte-autonomous effects on lipogenesis or lipid uptake. To analyze hepatocyte-specific functions of LSD1, we deleted *Lsd1* in hepatocytes of 6-week-old *db*/*db Lsd1^fl/fl^* mice using AAV8-TBG-iCre (*Lsd1^ΔL^ db*/*db* mice hereafter) (**Supplemental Figure 4B** and **C**). In contrast to systemic LSD1 inhibition, liver-specific *Lsd1* deletion did not alter body weight or blood glucose levels (**Supplemental Figure 4D** and **E**). Furthermore, glucose tolerance, plasma insulin levels, and insulin sensitivity were similar in *Lsd1^ΔL^ db*/*db* and control *db*/*db* mice (**Supplemental Figure 4F-H**). Accordingly, hepatocyte-specific *Lsd1* deletion did not improve pyruvate tolerance, reflecting unaltered hepatic gluconeogenesis (**Supplemental Figure 4I**). Further supporting this conclusion, LSD1 inhibition in primary hepatocytes isolated from *db*/*db* mice did not alter glucose production from lactate and pyruvate (**Supplemental Figure 4J**). Together, these findings suggest that improvements in glucose homeostasis following systemic LSD1 inhibition are not mediated by a direct effect on hepatocytes.

We further examined the effects of hepatocyte-specific *Lsd1* deletion on liver steatosis in *db/db* mice. *Lsd1^ΔL^ db*/*db* mice exhibited similar liver histology, liver weight, plasma triglycerides, ALT, and AST levels as *db/db* control mice (**Supplemental Figure 4K-O**). However, we observed a reduction in hepatic triglyceride levels in *Lsd1^ΔL^ db*/*db* mice (**Supplemental Figure 4P)**, suggesting a hepatocyte-autonomous role of LSD1 in the regulation of hepatic lipid metabolism. Despite the observed effect of hepatic *Lsd1* deletion on hepatic triglycerides, our findings support the overall conclusion that the beneficial effects of systemic LSD1 inhibition on glucose homeostasis and liver steatosis are not a direct result of LSD1 inhibition in hepatocytes.

### Food intake is reduced by systemic LSD1 inhibition

The improvements in glucose homeostasis and liver health following systemic LSD1 inhibition coincided with reductions in weight gain (**Figure 1B**). Changes in body weight typically result from imbalanced caloric intake and energy expenditure. To investigate whether GSK-LSD1-mediated effects on weight gain are caused by decreased food consumption or increased energy expenditure, GSK-LSD1-or veh-treated *db*/*db* mice were placed in metabolic cages and monitored over the course of 48 hours. This experiment revealed that food intake was reduced by GSK-LSD1 treatment (**Figure 4A** and **Supplemental Figure 5A**). In contrast, GSK-LSD1 had no significant impact on energy expenditure as measured by oxygen consumption and carbon dioxide production (**Figure 4B** and **C**), resulting in a similar respiratory quotient between GSK-LSD1- and veh-treated mice (**Supplemental Figure 5B**). We also found no evidence for changes in animal activity levels (**Supplemental Figure 5C**). Fluid intake was reduced by GSK-LSD1 (**Supplemental Figure 5D** and **E**), which is likely a secondary effect of normalized glucose levels. Importantly, long-term measurements of food intake revealed no difference between GSK-LSD1- or veh-treated lean *db*/+ mice (**Supplemental Figure 5F**). Overall, these findings suggest that systemic LSD1 inhibition reduces weight gain in *db/db* mice in part by attenuating hyperphagia.

**Figure 4.**
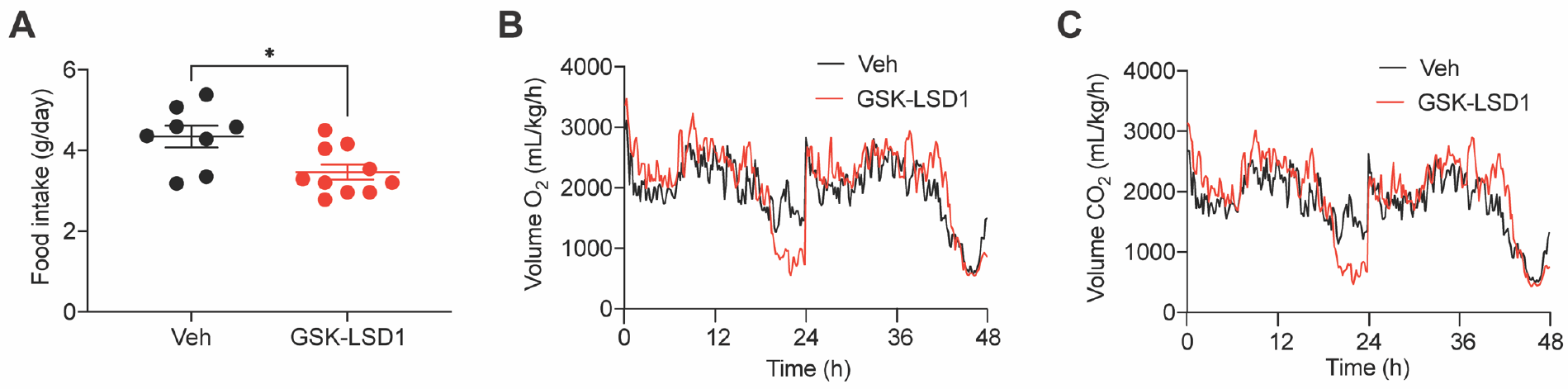
Food intake is reduced in GSK-LSD1-treated *db*/*db* mice. (**A**) Mice were injected daily with GSK-LSD1 or vehicle (veh) for 5 weeks and then placed into metabolic cages. Using the Comprehensive Laboratory Monitoring System (CLAMS), food intake was monitored over 48 hours and is shown as food consumed per day (n = 4-5 mice/group/day). (**B**) Oxygen consumption (V_O2_) and (**C**) carbon dioxide production (V_CO2_) (n = 4-5 mice/group). Data presented as mean ± SEM. Statistical differences between two groups were calculated using an unpaired two-tailed Student’s t-test (A) or a two-way ANOVA (B, C) with Tukey post hoc analysis. \**p*<0.05.

### Metabolic effects of systemic LSD1 inhibition are independent of food intake

Pair-feeding of *db/db* mice to lean controls has been shown to prevent obesity and hyperglycemia (38). Having observed reduced food intake and weight gain in GSK-LSD1-treated mice, we reasoned that some of the metabolic benefits of systemic LSD1 inhibition could result from its effect on feeding. Therefore, we conducted a pair-feeding experiment in which food consumption in GSK-LSD1-treated *db*/*db* mice was measured daily and the same amount was then provided to a second group of *db*/*db* mice receiving veh, with a third group of veh-treated ad libitum-fed *db*/*db* mice serving as a control (**Figure 5A**). GSK-LSD1-treated mice consumed significantly less food over the course of the study compared to veh-treated ad libitum-fed mice, whereas the veh-treated pair-fed group consumed the same amount of food as the GSK-LSD1-treated group as defined per experimental design (**Figure 5B** and **Supplemental Figure 6A**). Body weight gain did not differ between the three groups over the course of the intervention (**Supplemental Figure 6B**). Blood glucose levels in GSK-LSD1-treated *db*/*db* mice were significantly lower than in veh-treated pair-fed *db*/*db* mice (**Figure 5C**). Likewise, GSK-LSD1-treated *db*/*db* mice showed improved glucose tolerance relative to veh-treated pair-fed *db*/*db* mice (**Figure 5D**), indicating that GSK-LSD1 prevents hyperglycemia in *db*/*db* mice independent of food intake. Accordingly, fasting plasma insulin levels remained high in pair-fed compared to ad libitum-fed mice, but were significantly lower in the GSK-LSD1-treated group (**Figure 5E**). Together, these observations suggest that GSK-LSD1 improves insulin sensitivity in *db*/*db* mice independent of its effect on food intake.

**Figure 5.**
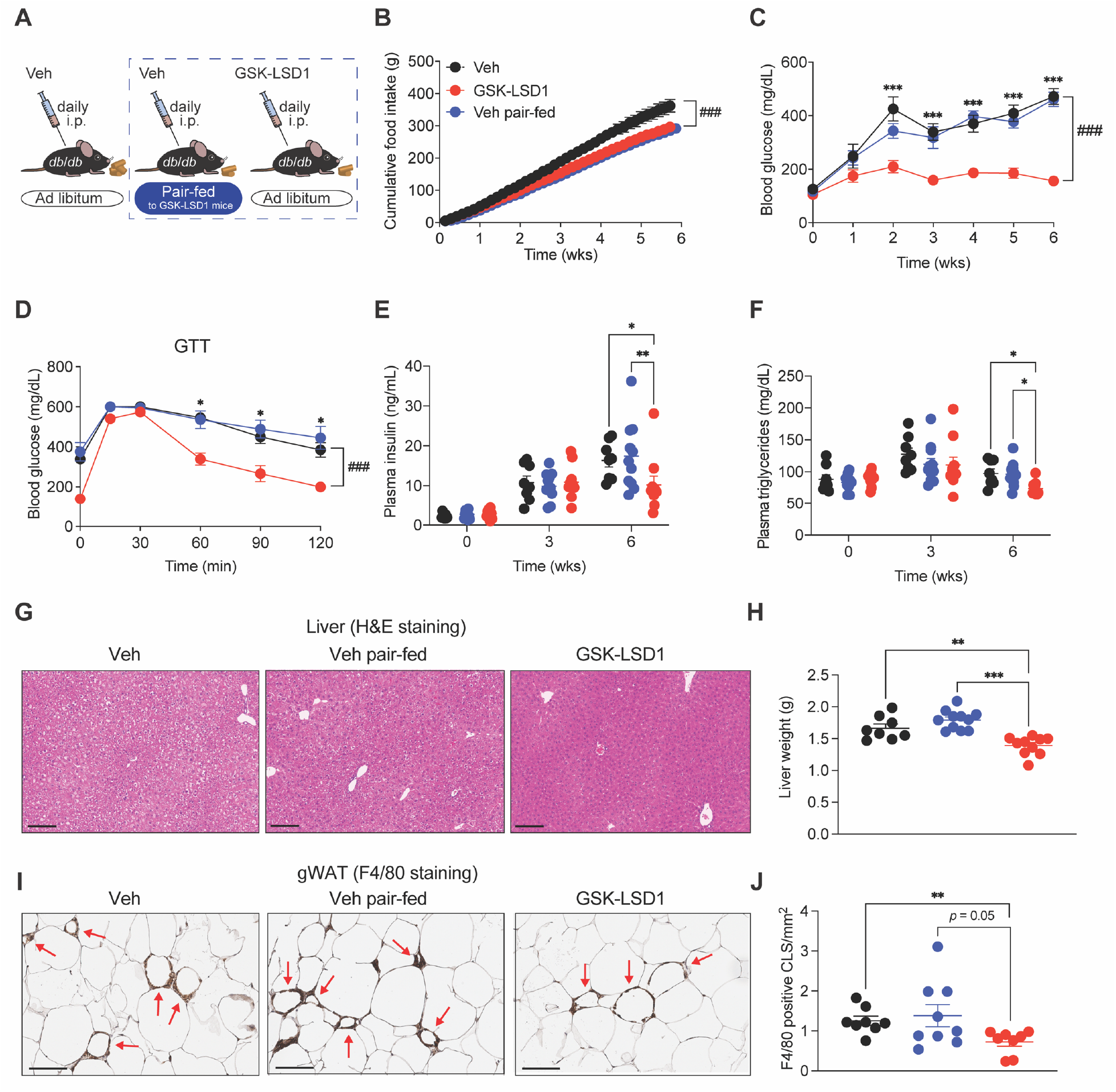
GSK-LSD1-mediated improvement of metabolic dysfunction is independent of reduced food intake. (**A**) Study design of pair-feeding experiment. 4-week-old *db*/*db* mice were injected daily with vehicle (veh) or GSK-LSD1 for 6 weeks and fed a normal chow diet ad libitum. A third group of mice received veh and was pair-fed to GSK-LSD1-treated mice. (**B**) Food consumption was monitored daily and is shown as cumulative food intake over the 6-week study (n = 8-11 mice/group). (**C**) Blood glucose levels measured weekly. Asterisks indicate statistical differences between veh (black) and GSK-LSD1 group (red). (**D**) Blood glucose levels at indicated time points after a glucose bolus via oral gavage (n = 8-11 mice/group). Glucose tolerance test, GTT. (**E**) Fasting plasma insulin levels at the indicated time points (n = 8-11 mice/group). (**F**) Fasting plasma triglycerides levels at indicated time points (n = 8-11 mice/group). (**G**) Representative images of liver sections stained with hematoxylin and eosin (H&E). Scale bars = 100 μm (n = 4 mice/group). (**H**) Liver weight (n = 8-11 mice/group). (**I**) Representative images of gWAT sections stained against F4/80 to detect crown-like structures (CLS). Red arrows indicate CLS. Scale bars = 100 μm. (**J**) Quantification of F4/80^+^ CLS in gWAT relative to tissue size (n = 8-9 mice/group). Data presented as mean ± SEM. Statistical differences were calculated using a one-way ANOVA (D, F, H, J) or two-way ANOVA (B, C, E) with Tukey post hoc analysis. \**p*<0.05, \*\**p*<0.01, \*\*\**p*<0.001, ^###^*p*<0.001, ns, not significant.

The feeding-independent effects of GSK-LSD1 upon glucose homeostasis suggested that systemic LSD1 inhibition could ameliorate complications of obesity independent of altered food intake. Indeed, LSD1 inhibition reduced plasma triglycerides, improved liver steatosis, reduced liver weight and plasma ALT activity compared to veh-treated pair-fed *db*/*db* mice (**Figure 5F-H** and **Supplemental Figure 6C**), indicating that these effects are independent of reduced feeding. Similarly, the effect of LSD1 inhibition on adipose tissue inflammation was independent of reduced food intake, as evidenced by reduced abundance of F4/80^+^ CLS compared to veh-treated pair-fed *db*/*db* mice (**Figure 5I** and **J**). Overall, these experiments show that beneficial effects of GSK-LSD1 on glucose homeostasis, liver health, and adipose inflammation occur independent of its hyperphagia-reducing effect.

### GSK-LSD1 prevents obesity and improves insulin signaling in Western diet-fed mice

*db/db* mice are deficient for signaling by the satiety hormone leptin and therefore do not fully recapitulate human obesity, in which leptin function is intact. Therefore, we determined whether effects of GSK-LSD1 can be observed in mice that develop obesity following consumption of a high-energy Western diet (39). C57BL/6J mice were fed a Western diet (42% kcal from fat, 34% sucrose by weight) for 11 weeks concomitant with daily injections of GSK-LSD1 or veh, with veh-treated C57BL/6J mice fed a normal chow diet serving as a control (**Figure 6A**). Western diet feeding resulted in substantial weight gain of veh-treated mice, while GSK-LSD1-treated mice fed the Western diet remained lean, exhibiting body weights similar to those of normal chow-fed mice (**Figure 6B**). Notably, GSK-LSD1 also prevented obesity in response to consumption of a high fat diet (HFD, 60% kcal from fat; **Supplemental Figure 7A**). While blood glucose levels did not change in response to either diet (**Figure 6C** and **Supplemental Figure 7B**), GSK-LSD1 improved glucose tolerance in mice fed a high fat diet but not a chow diet (**Supplemental Figure 7C**). Plasma insulin levels increased in response to Western diet feeding of veh-treated mice but remained low in the GSK-LSD1 group (**Figure 6D**). The reduction in insulin levels without corresponding increases of blood glucose in GSK-LSD1-treated mice suggests that LSD1 inhibition improves insulin sensitivity during Western diet feeding, as we observed in *db/db* mice.

**Figure 6.**
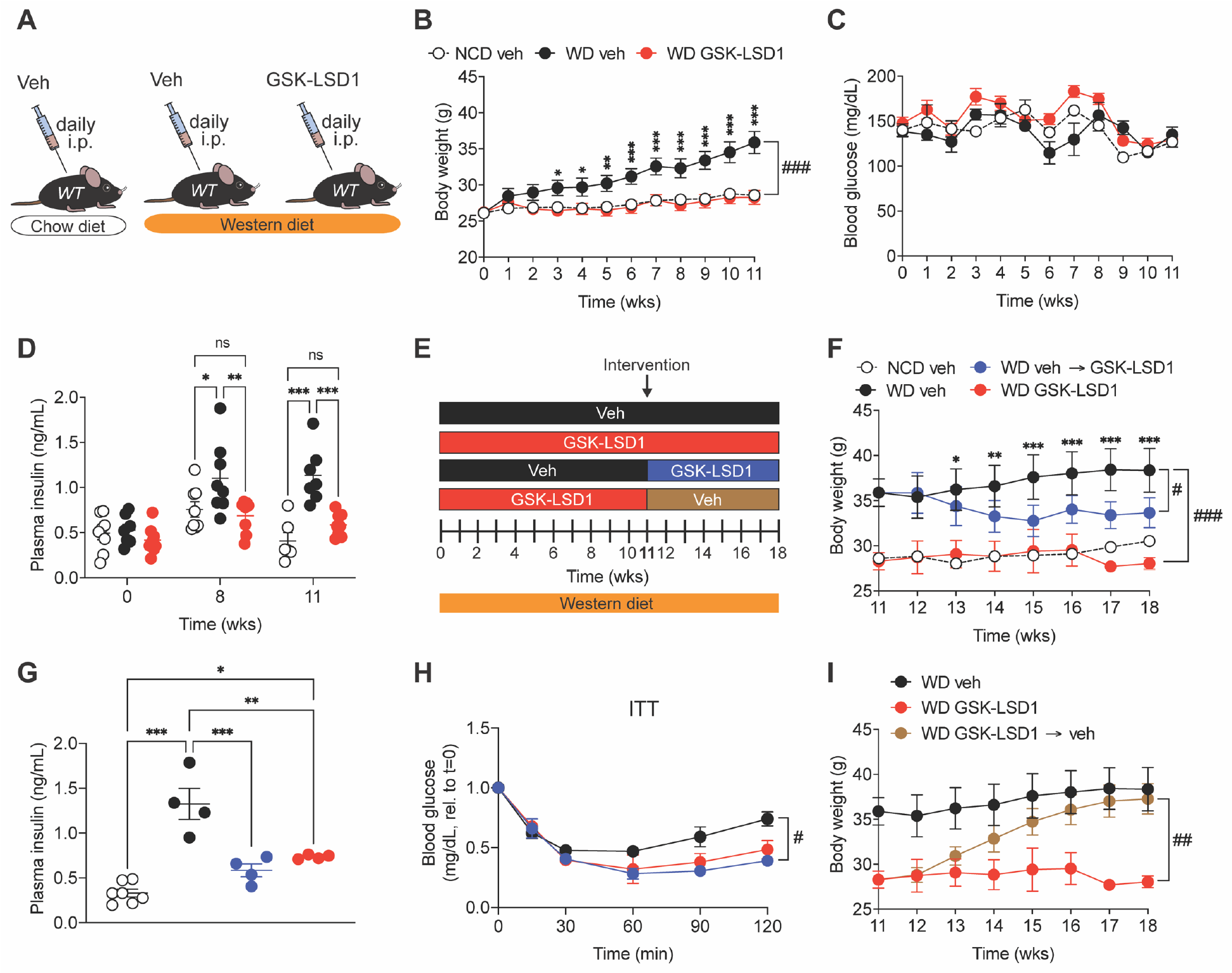
GSK-LSD1 reverses metabolic dysfunction due to diet-induced obesity. (**A**) 10-week-old C57BL/6J *wild type* (*WT*) mice fed a Western diet (WD) were injected daily with GSK-LSD1 or vehicle (veh) for 11 weeks. As a control, a group of *WT* mice received veh and was kept on a normal chow diet (NCD). (**B**) Body weight and (**C**) blood glucose levels measured weekly (n = 7-8 mice/group). Asterisks indicate statistical differences between GSK-LSD1- and veh-treated mice on WD unless otherwise stated. (**D**) Fasting plasma insulin levels at indicated time points (n = 6-8 mice/group). (**E**) Overview of intervention treatment protocol. After 11 weeks of veh administration, the veh group was split into one group continuing veh administration (black), whereas the other half began to receive GSK-LSD1 daily for another 7 weeks (blue). Likewise, the GSK-LSD1 group was split into one group continuing GSK-LSD1 administration after 11 weeks (red), whereas the other half began to receive veh daily for another 7 weeks (brown). (**F**) Body weight across all treatment groups (WD: n = 4 mice/group for week 11-16, n = 2-4 mice/group for week 17 and 18, n = 7 mice for NCD group). Asterisks indicate statistical differences between WD-fed mice administered veh (black) and the intervention group (blue). (**G**) Fasting plasma insulin levels 2 weeks after drug intervention (NCD veh: n = 7 mice, WD groups: n = 4 mice). (**H**) Blood glucose levels at indicated time points after intraperitoneal insulin (0.8 U/kg body weight) injection at week 17. Data shown relative to time point 0 min (n = 2-4 mice/group). Insulin tolerance test, ITT. (**I**) Body weight (WD: n = 4 mice/group for week 11-16, n = 2-4 mice/group for week 17 and 18). Data are shown as mean ± SEM. Statistical differences were calculated using a one-way ANOVA (G, J) or two-way ANOVA (B-D, F, H, I) with Tukey post hoc analysis. \**p*<0.05, \*\**p*<0.01, \*\*\**p*<0.001, ^#^*p*<0.05, ^##^*p*<0.01, ^###^*p*<0.001, ns, not significant.

The above findings collectively demonstrate that GSK-LSD1 prevents metabolic defects in several models of obesity. However, an ideal therapy for metabolic disease would reverse preexisting defects. Therefore, we performed a drug intervention study to ask whether GSK-LSD1 improves metabolic health in obese mice (**Figure 6E**). After 11 weeks of Western diet feeding and veh treatment, mice were split into two groups receiving either GSK-LSD1 for an additional 7 weeks or continuation of veh treatment. Remarkably, switching treatment groups from veh to GSK-LSD1 lowered body weight and plasma insulin levels significantly in Western diet-fed mice (**Figure 6F** and **G**). As blood glucose levels did not change in any of the treatment groups (**Supplemental Figure 8A**), this result suggests that GSK-LSD1 ameliorates insulin resistance in mice with preexisting obesity. Indeed, insulin tolerance tests revealed that insulin sensitivity was improved to a similar extent whether GSK-LSD1 was administered concomitant with Western diet feeding or initiated after the establishment of obesity (**Figure 6H**). Altogether, these observations support the potential for systemic LSD1 inhibition as a therapy for obesity and T2D.

GSK-LSD1 is an irreversible inhibitor, which could limit the ability to reverse the effects of this drug following treatment withdrawal (e.g., when target weight is achieved in previously obese individuals). To test whether the effects of GSK-LSD1 upon systemic metabolism are reversible, we performed a washout study in Western diet-fed mice treated with GSK-LSD1 for 11 weeks, at which point mice were switched to veh treatment (**Figure 6E**). Western diet-fed mice that received GSK-LSD1 or veh for the duration of the study were used for comparison. The GSK-LSD1-mediated prevention of Western diet-induced body weight gain was completely reversible once the mice were taken off treatment, with body weights of the group taken off GSK-LSD1 converging with those of mice treated with veh for the duration of the study (**Figure 6I**). Similarly, beneficial effects of GSK-LSD1 on liver, gWAT, and sWAT weight were reversed when Western diet-fed mice were taken off the drug (**Supplemental Figure 8B-D**). Overall, our findings indicate that systemic LSD1 inhibition prevents and corrects hallmark metabolic defects associated with obesity in a manner that is readily reversible upon drug withdrawal.

## Discussion

Despite the growing obesity pandemic, there is an unmet need for efficient and safe therapies for obesity and its associated complications. Here, we report that systemic LSD1 inhibition in rodent models of obesity and T2D causes weight loss and ameliorates obesity-induced complications including hyperglycemia and NAFLD. LSD1 inhibition decreases food intake, body weight, and fat mass in diet-induced and genetic models of obesity. Moreover, GSK-LSD1 prevents hyperglycemia and fatty liver disease in *db*/*db* mice. These effects are accompanied by improved adipose tissue function as shown by reduced tissue inflammation and increased insulin sensitivity. LSD1 inhibition does not affect metabolism in lean mice, indicating LSD1 plays a context-specific role during overfeeding. This is of importance for clinical applications, as LSD1 inhibition does not cause anorexia or hypoglycemia, suggesting physiological set points are maintained during LSD1 inhibition. Overall, these findings identify LSD1 as a potential therapeutic target to promote weight loss and prevent T2D and NAFLD in people with obesity.

The beneficial effects of LSD1 inhibition on metabolism occurred independent of reduced food intake, suggesting direct effects upon tissues involved in nutrient metabolism. Our findings indicate that LSD1 inhibition in WAT has a major contribution to the beneficial effect of GSK-LSD1 on obesity-associated metabolic complications. In support of this conclusion, improved systemic insulin sensitivity after GSK-LSD1 treatment was associated with improved insulin signaling in WAT but not in skeletal muscle or liver. In addition, GSK-LSD1 reversed several obesity-associated defects in adipose that are known to contribute to metabolic syndrome, including inflammation and excessive lipolysis. LSD1 has been reported to have a cell-autonomous role in hepatocytes, promoting steatosis through transcriptional activation of lipogenic enzyme genes (29). Consistent with this finding, we observed lower hepatic triglyceride levels after hepatocyte-specific *Lsd1* deletion. However, we found that hepatocyte-specific *Lsd1* deletion did not recapitulate the beneficial effect of systemic GSK-LSD1 treatment upon NAFLD or hepatic glucose production, suggesting that systemic LSD1 inhibition indirectly ameliorates hepatic steatosis and excessive gluconeogenesis.

It is well established that obesity-associated changes in adipose tissue can have pleiotropic effects on systemic metabolism (1, 4, 37, 40). Excess caloric intake and the subsequent expansion of fat mass result in increased infiltration of macrophages and local secretion of proinflammatory cytokines, which precipitate metabolic defects in adipocytes that lead to increased circulating FFA (4, 37). This sets in motion a vicious cycle of additional macrophage recruitment, further inflammation, lipolysis, and insulin resistance (1, 5, 6, 37, 40–46). Our finding that GSK-LSD1 inhibits lipolysis in isolated adipocytes indicates effects of LSD1 on lipolysis independent of inflammation and insulin signaling, suggesting a direct effect of LSD1 on lipolysis could initiate a chain of events leading to adipose tissue inflammation and insulin resistance in vivo. In support of this model, inhibition of lipolysis has been shown to improve insulin sensitivity, glucose tolerance, adipose tissue inflammation, and liver health during obesity (37, 47). In addition to the GSK-LSD1-mediated reduction in lipolysis in isolated adipocytes, we also found decreased circulating FFA levels in vivo. It is known that fatty acids released from adipose promote hepatic glucose production and are the main substrate for hepatocyte triglyceride synthesis (5, 44, 48–52). Consequently, lipolysis-derived FFA promote hepatic gluconeogenesis and ectopic lipid deposition in hepatocytes (37, 47), providing a plausible link between lipolysis inhibition by GSK-LSD1 and indirect metabolic improvements in the liver. Further studies will be necessary to clarify the interrelationship between lipolysis inhibition, reduced adipose inflammation, and improved insulin signaling following systemic LSD1 inhibition.

Genetic studies in lean mice have revealed that a major function of LSD1 is to promote mitochondrial metabolism in adipocytes (33, 53). *Lsd1* deletion in white and brown fat via adiponectin-Cre leads to increased adipose tissue mass and weight gain (53). Moreover, this model develops exaggerated glucose intolerance during HFD feeding compared to *Lsd1*-intact controls (53). Our finding that pharmacological LSD1 inactivation reduces adipose tissue mass and improves glucose tolerance during obesity seemingly contrasts with these genetic studies. It is possible that pharmacological inhibition of LSD1’s enzymatic activity has a distinct effect from genetic deletion of *Lsd1*, which can impact other functions such as scaffolding of proteins within transcriptional complexes. Alternatively, the use of constitutive Cre recombinase to delete *Lsd1* may have confounding effects on adipocyte development (33).

The selective effect of LSD1 inhibition during overfeeding suggests LSD1 has context-dependent functions in the regulation of systemic metabolism. We found that GSK-LSD1 reduces food intake and body weight only in models of genetic or diet-induced obesity, with no effect on these parameters in lean, chow-fed mice. Similarly, LSD1 inhibition prevents hyperglycemia in *db*/*db* mice but is of little effect on glycemia or glucose tolerance in lean, nondiabetic mice. Obesity and T2D are known to disrupt homeostatic feedback mechanisms regulating appetite, weight gain, and blood glucose (13). Our observations open the possibility that LSD1 is involved in the maladaptive changes to systemic metabolism that result in higher defended body weight and blood glucose. Effects of LSD1 inhibition on food intake, adipose lipolysis, hepatic triglycerides, and gluconeogenesis suggest LSD1 could evoke shared tissue-autonomous mechanisms for metabolic rewiring during obesity. While our pharmacological approach does not definitively implicate a single target tissue responsible for these effects, simultaneously targeting several relevant tissues by GSK-LSD1 treatment provides the advantage of testing the coordinated effects of LSD1. Future work should address whether and how obesity-associated changes to LSD1 and its associated epigenomic program occur in endocrine and neuronal systems involved in feeding behavior as well as in peripheral tissues directly involved in fuel metabolism.

The development of weight loss therapies to treat obesity and its complications has proven to be a considerable challenge. None of the obesity drugs with longstanding FDA approval are widely used due to the risk of adverse events (54, 55). The recently approved glucagon-like peptide 1 receptor agonist semaglutide holds promise for safe and effective obesity treatment when combined with lifestyle interventions (56). With the hope that broad adoption of semaglutide mirrors results of clinical trials indicating durable weight loss and reduced complications of obesity (56, 57), the next generation of obesity drugs may well be tailored to patients at risk for specific complications such as NAFLD, for which there is currently no approved therapy beyond implementing lifestyle changes. Epigenomic regulation has recently emerged as a novel regulatory layer in energy homeostasis, opening a new avenue for potential treatment strategies for metabolic disease (58, 59). Evidence that histone deacetylases (HDACs) play a role in glucose homeostasis by regulating the function of hepatocytes and insulin-producing pancreatic β-cells positioned these enzymes as potential drug targets in metabolic disease (58). Multiple HDAC inhibitors are currently being investigated for the treatment of obesity and T2D (58), but currently none have been approved for this clinical application. Here, we show that inhibition of the histone demethylase LSD1 has beneficial effects on glucose homeostasis and NAFLD independent of changes in food intake, suggesting that LSD1 could be a novel therapeutic target for correcting epigenomic defects in metabolic disease. The feasibility of LSD1 targeting in people with obesity will depend upon a favorable safety profile. Our results describing metabolic benefits of GSK-LSD1 provide a strong rationale for clinical trials investigating the safety and efficacy of LSD1 inhibitors in obesity and NAFLD.

## Methods

### Mice

Mice homozygous for the leptin receptor (*BKS.Cg-Dock7^m^ +/+ Lepr^db^/J*, JAX #000642 thereafter referred to as *db*/*db*) and the respective lean control animals (*db*/+) as well as C57BL/6J mice (JAX #000664) were purchased from The Jackson Laboratory (60, 61). *Lsd1*^fl/fl^ mice (provided by the laboratory of Michael Rosenfeld) (62) were bred with *db*/*db* mice to generate *Lsd1*^fl/fl^ *db*/*db* mice. Liver-specific *Lsd1* deletion was induced by intravenous injection of AAV8-TBG-iCre virus (2.5 x 10^11^ genome copies/mouse). As controls, *Lsd1*^fl/+^*db*/*db* and *Lsd1*^+/+^*db*/*db* mice were injected with the virus in parallel. All mice were housed and bred in vivaria approved by the Association for Assessment and Accreditation of Laboratory Animal Care located in the School of Medicine, UCSD, following standards and procedures approved by the UCSD Institutional Animal Care and Use Committee. Mice were weaned at 4 weeks, maintained on a 12-hour light cycle, and fed ad libitum with water and standard rodent chow (PicoLab® Rodent Diet 20 5053), a Western diet (TD.88137 Envigo Teklad) containing 42 % kcal from fat, or a high fat diet (HFD, D12492 Research Diets) containing 60 % kcal from fat, unless otherwise indicated. Both male and female mice were used for all studies on *db*/*db* mice, male mice only were used for Western diet and HFD studies. Mice received GSK-LSD1 (500 μg/kg/mouse, Sigma SML1072) or vehicle (0.9% NaCl) daily via intraperitoneal injections. Body weight and blood glucose levels were monitored weekly.

### Metabolic studies

For glucose tolerance tests (GTT), mice were fasted for 5 h and orally gavaged with 2 mg/g body weight glucose. Plasma glucose levels were measured in blood samples from the tail vein at baseline, 15 min, 30 min, 60 min, 90 min, and 120 min after gavage using a Bayer Contour glucometer (Bayer). For insulin tolerance tests (ITT), an insulin solution (0.8 – 2.0 U/kg body weight) was intraperitoneally injected into fasted (5 h) mice and glucose levels were monitored as described above. For pyruvate tolerance tests (PTT), overnight starved mice (16 h) were intraperitoneally injected with a pyruvate solution (1 mg/g body weight) and plasma glucose levels were analyzed as described above.

### ELISA

Plasma insulin levels were measured after 5 h of fasting or before and 10 min after a glucose gavage (2 mg/g body weight) of fasted (5 h) mice via the mouse ultrasensitive or mouse insulin ELISA kit (Alpco).

### Acute insulin response

To determine tissues insulin responsiveness, mice were starved overnight (16 h) and intraperitoneally injected with 2.0 U/kg body weight of an insulin solution. Tissues were harvested 15 min after insulin stimulation and flash frozen in liquid N_2_. Lysates were then prepared of gWAT, liver, and skeletal muscle and samples were subsequently analyzed via Western blotting.

To measure insulin sensitivity in vitro, differentiated adipocytes were pre-incubated with GSK-LSD1 or veh overnight. The next day, cells were serum starved in Krebs buffer for 2 h and stimulated with 10 mM Insulin for 10 min. Treatment with GSK-LSD1 or veh was continued throughout the experiment. Adipocytes were then washed with ice cold PBS, lysed in RIPA buffer (Thermo Fisher) containing protease inhibitor and phosphatase inhibitor and protein content was determined using commercial kits. Insulin responsiveness in adipocytes was then quantified by visualizing Akt and p-Akt via Western blotting.

### Western blotting

Tissues were homogenized using RIPA buffer (Thermo Fisher Scientific) and freshly added protease inhibitor cocktail (Roche) and phosphatase inhibitor (Sigma). Lysates of gWAT (20 µg), liver (40 µg), and skeletal muscle (40 µg) of *db*/*db* mice and of liver (30 µg) from *Lsd1^Δ^*^L^*db*/*db* mice were analyzed by SDS-PAGE on 12% Bis-Tris gels with equal amount of protein loading. Proteins were visualized after transfer onto a PVDF blotting membrane (GE Healthcare) and incubation with specific primary antibodies using horseradish peroxidase-conjugated secondary antibodies. Western blot primary antibodies included: rabbit anti-mouse Akt (Cell Signaling Technology, #4691S), rabbit anti-mouse p-Akt Ser473 (Cell Signaling Technology, #4060S), mouse anti-mouse GAPDH (Thermo Fisher, #AM4300), rabbit-anti-mouse LSD1 (Abcam, #17721) and mouse anti-mouse Vinculin (Abcam, #ab18058). Western blot secondary antibodies included: goat anti-rabbit HRP (Southern Biotech, #4010-05) and ECL sheep anti-mouse (GE Healthcare #NA931V).

### Histology

Immunohistochemical analysis were performed on sections of paraformaldehyde-fixed and paraffin-embedded tissues. Histology was performed by the University of California San Diego Histology and Immunohistochemistry core. Routine hematoxylin and eosin (H&E) stain was performed on adipose tissues, liver, and skeletal muscle. The size of adipocytes was determined by quantifying at least 100 adipocytes using ImageJ analysis software (NIH).

To analyze crown-like structures, adipose tissues were stained with rat anti-F4/80 primary antibody (Bio-Rad, #MCA497B) and goat anti-rat HRP polymer (Cell IDX, #AH-100) followed by chromogenic 3,3’-diaminobenzidine (VWR, #95041-478) and counter stained with val hematoxylin (Biocare, #VLT8014G20). Slides were imaged using a slide scanner machine (NanoZoomer, Hamamatsu) and blinded samples were analyzed using ImageJ analysis software (NIH). The number of crown-like structures was quantified on whole tissue sections.

### RNA analysis

Total RNA was isolated in Trizol from homogenized tissue and cells and purified using RNeasy columns and Rnase free Dnase digestion according to the manufacturer’s instructions (QIAGEN). The quality and quantity of the total RNA was monitored and measured with a NanoDrop (NanoDrop Technologies, Inc. Wilmington, DE) following the manufacturer’s instructions. For quantitative PCR analysis, cDNA was synthesized using the iScript™ cDNA Synthesis Kit (Bio-Rad) and 500 ng of isolated RNA per reaction. 20 ng of template cDNA per reaction and iQ™ SYBR^®^ Green Supermix (Bio-Rad) were used for real-time PCR with gene-specific primers (Supplemental Table 1) and *Tbp* as a house keeping gene on a CFX96™ Real-Time PCR Detection System (Bio-Rad).

### Lipolysis in primary adipocytes

Lipolysis was measured in differentiated adipocytes derived from the stromal vascular fraction of subcutaneous white adipose tissue (sWAT). In detail, sWAT harvested from 5-week-old *db*/*db* mice was minced and digested with collagenase D (2.5 mg/mL, Sigma) for 30 min shaking at 37°C. The digestion was stopped by adding 10 mL culture medium (DMEM/F12 containing 10% FBS, 100 units/ml penicillin, and 0.1 mg/ml streptomycin) and the homogenate was filtered through a 100 µm cell strainer. After a 5 min centrifugation at 500 x g, the pellet containing the stromal vascular fraction was collected and incubated with erythrocyte lysis buffer for 5 min at room temperature. Next, the stromal vascular fraction was precipitated, and the cell pellet was then resuspended in culture medium and seeded into a T75 flask.

Once the pre-adipocytes reached 80% confluency, the cells were seeded into 12-wells and grown to confluency. Adipocyte differentiation was induced by adding 0.1 µM dexamethasone, 450 µM isobutylmethylxanthine, 2 µg/mL insulin, and 1 µM rosiglitazone (day 0). After 3 days, the medium was changed to culture medium containing 2 µg/mL insulin and 1 µM rosiglitazone. On day 5, the medium was changed to culture medium and the adipocytes were grown for another two days.

Differentiated adipocytes were pre-incubated with 1 µM GSK-LSD1 or veh overnight. The next day (day 7), adipocytes were serum starved in DMEM containing 2% fatty acid free BSA supplemented with 10 µM isoproterenol or control. After 1 h, the incubation media was replaced, and the release of fatty acids (Wako chemicals) and glycerol (Sigma) was determined after 4h. Treatment with GSK-LSD1 or veh was continued throughout the experiment. Data were normalized to cellular protein content as described above.

### Lipid and metabolic analysis

Lipid levels were analyzed in plasma and liver samples as previously described (63, 64). Blood was drawn via the tail vein from mice fasted for 5 h. Total cholesterol and triglyceride levels were determined in plasma and liver extracts using commercially available kits (Sekisui Diagnostics). Further, plasma β-hydroxybutyrate (Cayman Chemical), alanine aminotransferase and aspartate aminotransferase (Sigma) levels were determined using enzymatic kits. NEFA (Wako Chemicals) was measured in plasma obtained from overnight fasted mice.

### Metabolic cages

To determine energy expenditure and food intake, *db*/*db* mice administered with veh or GSK-LSD1 were placed into metabolic cages. Over a 48 h time frame, carbon dioxide production (VCO_2_), oxygen consumption (VO_2_), food intake, consumed water, and spontaneous motor activity were analyzed using the Comprehensive Laboratory Monitoring System (CLAMS, Columbus Instruments) at the University of California San Diego Animal Phenotyping Core. The respiratory quotient was calculated as the ratio of VCO_2_ to O_2_.

### Pair-feeding study

A pair-feeding experiment was performed on 4-week-old, individually housed *db*/*db* mice (both males and females), which were randomly assigned to the respective treatment groups. To establish pair-feeding conditions, a group of mice were administered with GSK-LSD1 as described above and fed a normal chow diet ad libitum. Food intake was measured daily between 4 and 6 pm and the average daily food consumption was calculated across all mice in this treatment group with separate calculations for male and female mice. Next, the vehicle treated pair-fed mice were provided with the calculated average amount of food consumed by the GSK-LSD1 treated mice over the last 24 h. This procedure was then continued daily for 6 weeks. Leftover food of the pair-fed group, if any, was weighted and removed from the cage before new food was added. As a control group, vehicle administered *db*/*db* mice were fed normal chow diet ad libitum for 6 weeks.

### Glucose production in primary hepatocytes

Primary hepatocytes were isolated from 5-week-old *db*/*db* mice by perfusion of the liver with EDTA to dissociate the cells, followed by Percoll centrifugation as described (65). Hepatocytes were seeded into collagen-coated 12-well plates at 300.000 cells/well, cultured in DMEM containing 10% FBS, 100 units/mL penicillin, and 0.1 mg/mL streptomycin and pre-incubated with 1 µM GSK-LSD1 or veh overnight. The next day, hepatocytes were serum starved in Krebs buffer (125 mM NaCl, 4 mM KCl, 1 mM CaCl_2_, 1 mM MgCl_2_, 0.85 mM KH_2_PO_4_, 1.25 mM Na_2_HPO_4_, 15 mM NaHCO_3_, 10 mM HEPES, 0.2% fatty acid free BSA) for 1h and gluconeogenesis was stimulated by adding 1 mM pyruvate and 10 mM lactate. After 2h, glucose production was measured using the HK assay kit (Sigma). Cells were lysed in RIPA buffer (Thermo Fisher) containing protease inhibitor and phosphatase inhibitor and protein content was determined using commercial kits.

### Statistics

Statistical analyses were performed using Graphpad Prism 8 (GraphPad Software). Normality was tested via Shapiro-Wilk test and F-tests were performed to analyze equal variances. Data that passed both tests were analyzed by two-tailed Student’s t-test for two-group comparisons and one-way ANOVA for comparison of multiple groups (> 2) followed by Tukey’s post hoc testing. For data with multiple variables, e.g. glucose measurements over time, a two-way ANOVA for repeated measurements followed by Tukey’s post hoc test or Fisher Least Significant Difference post hoc testing was performed. All data are presented as mean ± SEM. *P* values less than 0.05 were considered significant.

### Study approval

All mouse experiments were approved by the UCSD Institutional Animal Care and Use Committee (IACUC).

## Author contributions

B.R. designed research studies, conducted experiments, acquired data, analyzed data, and wrote the manuscript. D.P.P., H.Z., C.N., A.R.H., and I.O., conducted experiments and acquired data. P.L.S.M. designed research studies. M.W. designed research studies, conducted experiments, acquired data, analyzed data, and wrote the manuscript. M.S. designed research studies, analyzed data, and wrote the manuscript.

## Acknowledgments

We thank members of the Sander laboratory and Drs. Martin Myers and Jerrold Olefsky for helpful discussions. We thank Fenfen Liu, Vivian Lin, and Johanna Fleischman for technical assistance.

## Supplemental material

### Supplemental figures

**Figure S1.**
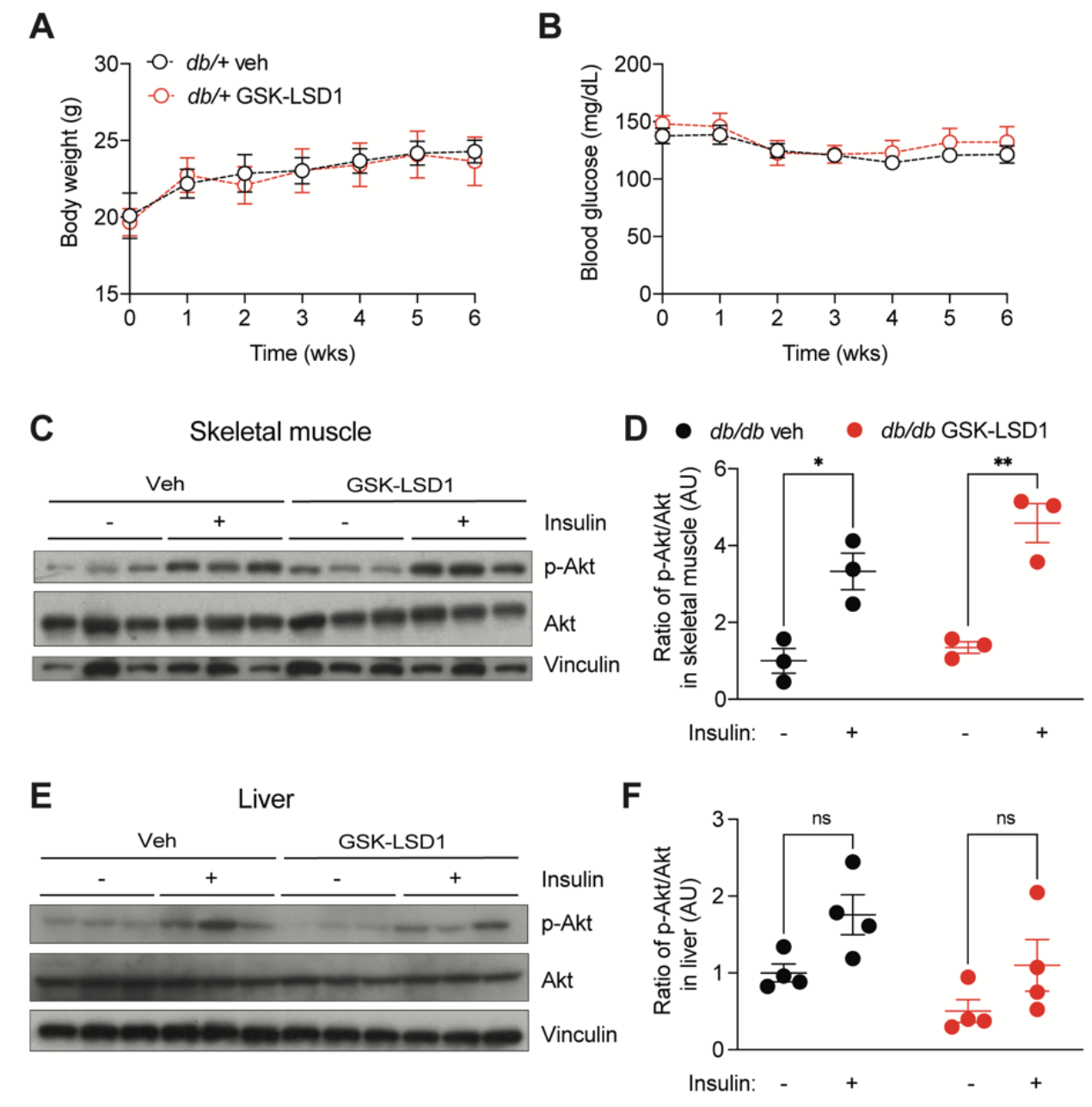
Impact of LSD1 inhibition on insulin sensitivity in peripheral tissues. (**A**) Body weight and (**B**) blood glucose levels in 4-week-old lean *db*/+ mice treated with GSK-LSD1 or vehicle (veh), respectively, for 6 weeks (n = 6 mice/group). (**C**-**F**) Immunoblot analysis of pAkt^Ser473^, Akt, and vinculin in (**C**) skeletal muscle (n = 3 mice/group) and (**E**) liver (n = 4 mice/group). Quantification of pAkt^Ser473^ to Akt ratio as fold change compared to veh-treated mice without insulin injection in skeletal muscle (**D**) and liver (**F**). Data are shown as mean ± SEM. Statistical differences were calculated using a two-way ANOVA with Tukey post hoc analysis. \**p*<0.05, \*\**p*<0.01. AU, arbitrary units, ns, not significant.

**Figure S2.**
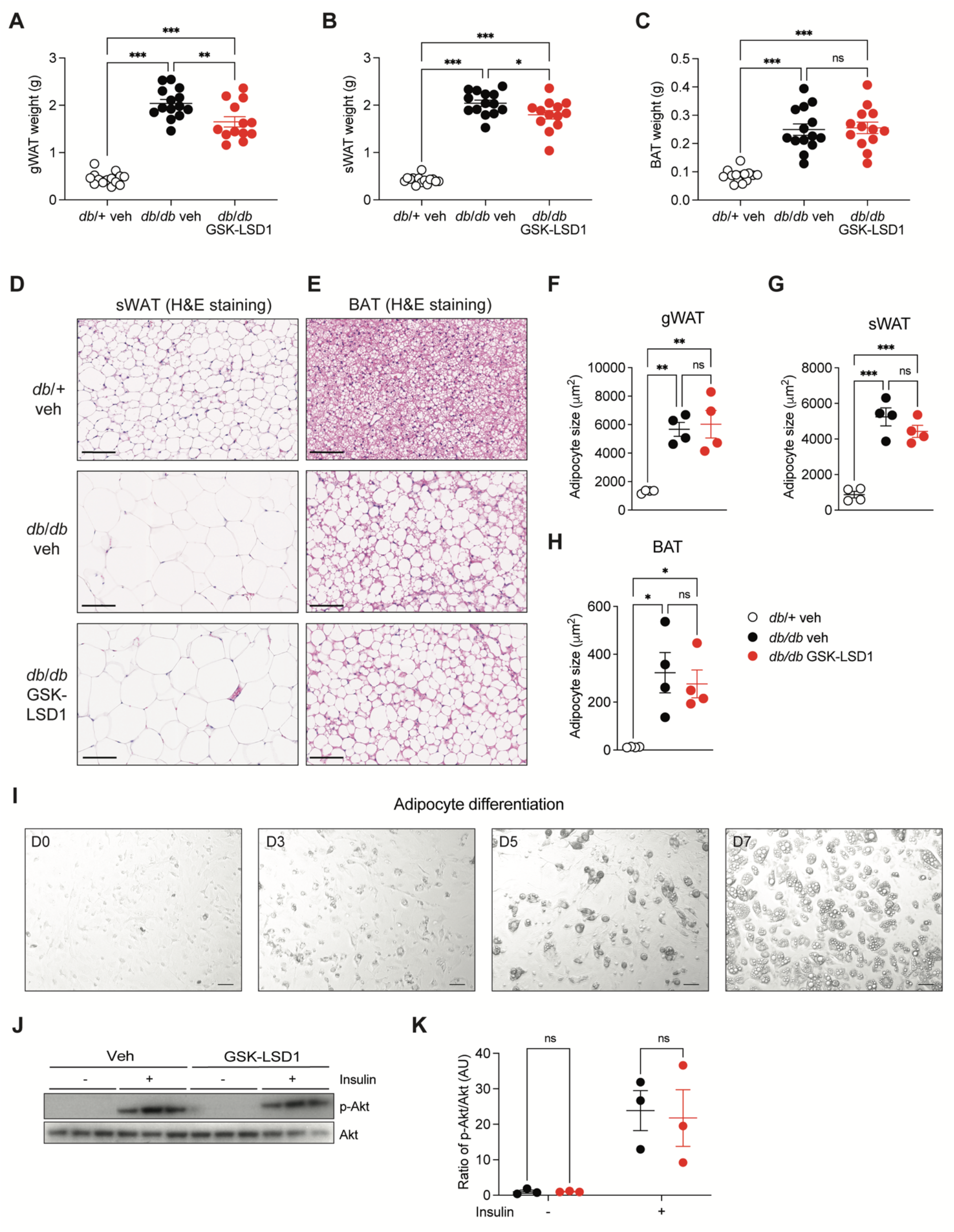
Impact of LSD1 inhibition on adipose tissue. (**A-H**) 4-week-old *db*/*db* mice received daily intraperitoneal (i.p.) injections of GSK-LSD1 or vehicle (veh) for 6 weeks. As a control, lean *db*/+ mice were injected with veh. Tissue weight of (**A**) gWAT, (**B**) sWAT, and (**C**) BAT (n = 13-14 mice/group). (**D**) Representative images of sWAT and (**E**) BAT sections stained with hematoxylin and eosin (H&E). Scale bars = 100 µm. (**F**) Adipocyte size quantified in gWAT, (**G**) sWAT, and (**H**) BAT (n = 4 mice/group). (**I**) Adipocytes were isolated from 5-week-old *db*/*db* mice. Representative images of adipocyte differentiation at day 0, 3, 5, and 7. Scale bars = 200 µm. (**J**) Immunoblot analysis of pAkt^Ser473^ and Akt in differentiated adipocytes isolated from *db*/*db* mice after preincubation with GSK-LSD1 or veh and stimulation with insulin (n = 3 mice). (**K**) Quantification of pAkt^Ser473^ to Akt ratio as fold change compared to veh-treated adipocytes. Data presented as mean ± SEM. A one-way ANOVA with Tukey post hoc analysis was performed to analyze statistical differences between three or more groups. \**p*<0.05, \*\**p*<0.01, \*\*\**p*<0.001, ns, not significant.

**Figure S3.**
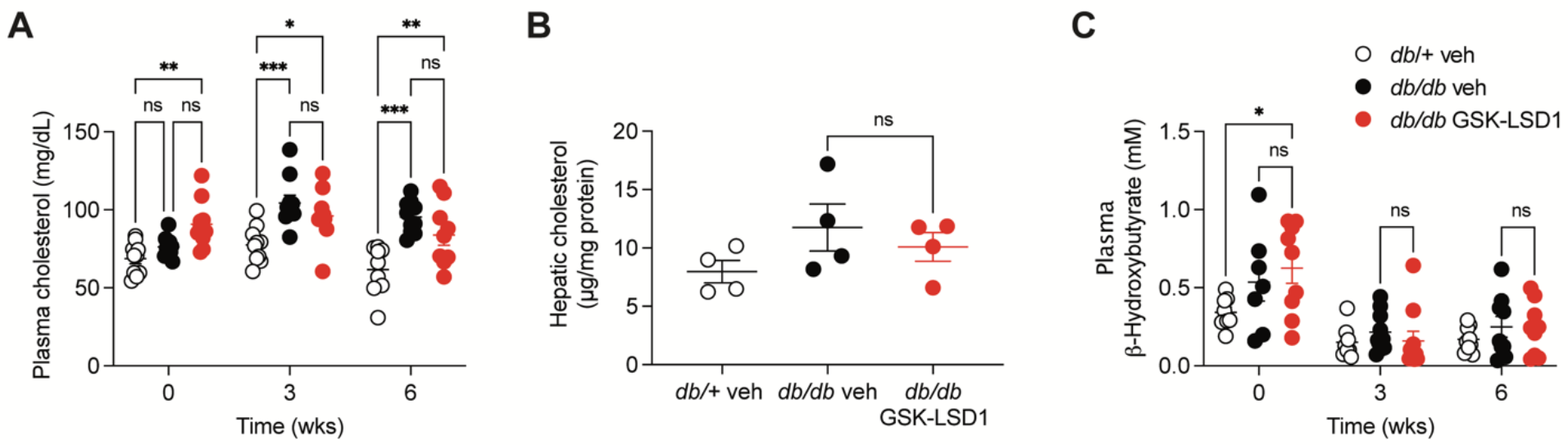
Plasma and hepatic metabolites in *db*/*db* mice. (**A**) 4-week-old *db*/*db* mice received daily intraperitoneal injections of GSK-LSD1 or vehicle (veh) for 6 weeks. As a control, lean *db*/+ mice were injected with veh. Fasting plasma cholesterol levels at indicated time points (n = 9-10 mice/group). (**B**) Hepatic cholesterol levels (n = 4 mice/group). (**C**) Plasma β-hydroxybutyrate levels at indicated time points (n = 7-10 mice/group). Data are shown as mean ± SEM. Statistical differences were calculated using a one-way ANOVA (B) or two-way ANOVA (A, C) with Tukey post hoc analysis. \**p*<0.05, \*\**p*<0.01, \*\*\**p*<0.001, ns, not significant.

**Figure S4:**
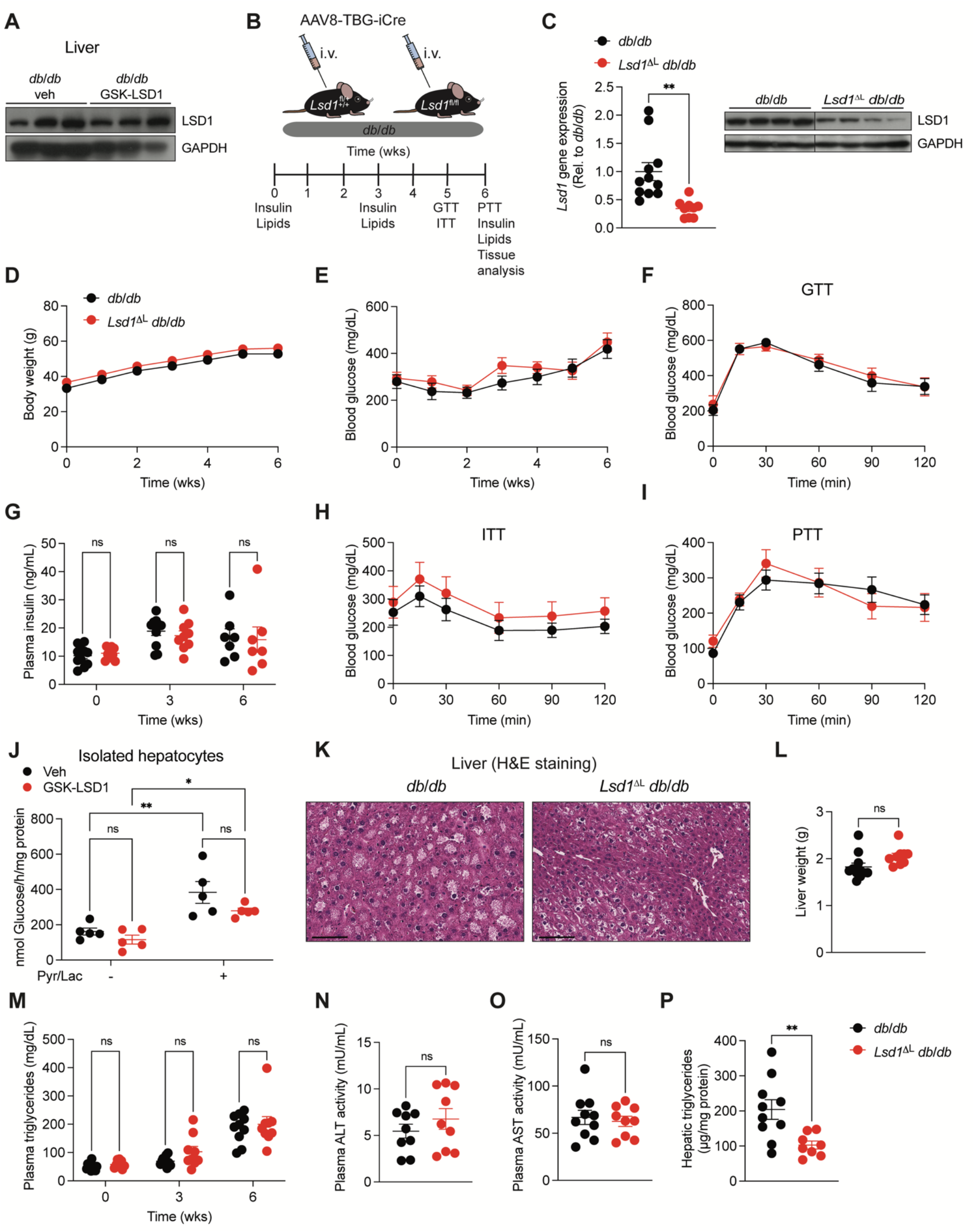
Hepatocyte-specific *Lsd1* deletion does not improve metabolic health in *db/db* mice. (**A**) Immunoblot analysis of LSD1 and GAPDH in the liver of GSK-LSD1- or vehicle (veh)-treated *db*/*db* mice (n = 3 mice/group). (**B**) 5-week-old *Lsd1*^fl/fl^*db*/*db* or control (*Lsd1*^fl/+^*db*/*db* and *Lsd1*^+/+^*db*/*db*) mice were intravenously injected with AAV8-TBG-iCre virus for hepatocyte-specific *Lsd1* deletion (hereafter referred to as *Lsd1^Δ^*^L^*db*/*db* mice). (**C**) qPCR analysis of *Lsd1* in livers of *Lsd1^Δ^*^L^*db*/*db* mice (left). Transcript levels relative to *db*/*db* control mice. (n = 9-11 mice/group). Immunoblot analysis of LSD1 and GAPDH in the liver of *Lsd1^Δ^*^L^*db*/*db* and control mice (right). The lanes were run on the same gel but were noncontiguous (n = 4 mice/group). (**D**) Body weight and (**E**) blood glucose levels measured weekly (n = 9-11 mice/group). (**F**) Blood glucose levels at indicated time points after a glucose bolus via oral gavage (n = 9-11 mice/group). Glucose tolerance test, GTT. (**G**) Fasting plasma insulin levels at baseline and after 3 and 6 weeks of GSK-LSD1 or veh treatment (n = 9-11 mice/group for week 0 and 3, n = 7 mice/group for week 6). (**H**) Blood glucose levels at indicated time points after intraperitoneal insulin (2.0 U/kg body weight) injection (n = 9-11 mice/group). Insulin tolerance test, ITT. (**I**) Blood glucose levels at indicated time points after intraperitoneal pyruvate injection (n = 9-11 mice/group). Pyruvate tolerance test, PTT. (**J**) Glucose production in hepatocytes isolated from 5-week-old *db*/*db* mice after preincubation with GSK-LSD1 or veh overnight. Glucose release into the medium was measured after 2 hours of pyruvate and lactate stimulation (n = 5 mice). (**K**) Representative images of liver sections stained with hematoxylin and eosin (H&E). Scale bars = 100 µm (n = 5-7 mice/group). (**L**) Liver weight (n = 9-11 mice/group). (**M**) Fasting plasma triglyceride levels at indicated time points. (**N**) Plasma ALT and (**O**) AST activity (n = 9-11 mice/group). (**P**) Hepatic triglyceride levels (n = 8-10 mice/group). Data are shown as mean ± SEM. Statistical differences were calculated using an unpaired Student’s t-test (L, O-P) or two-way ANOVA (D-J, M) with Tukey post hoc analysis. \**p*<0.05, \*\**p*<0.01, \*\*\**p*<0.001, ns, not significant.

**Figure S5.**
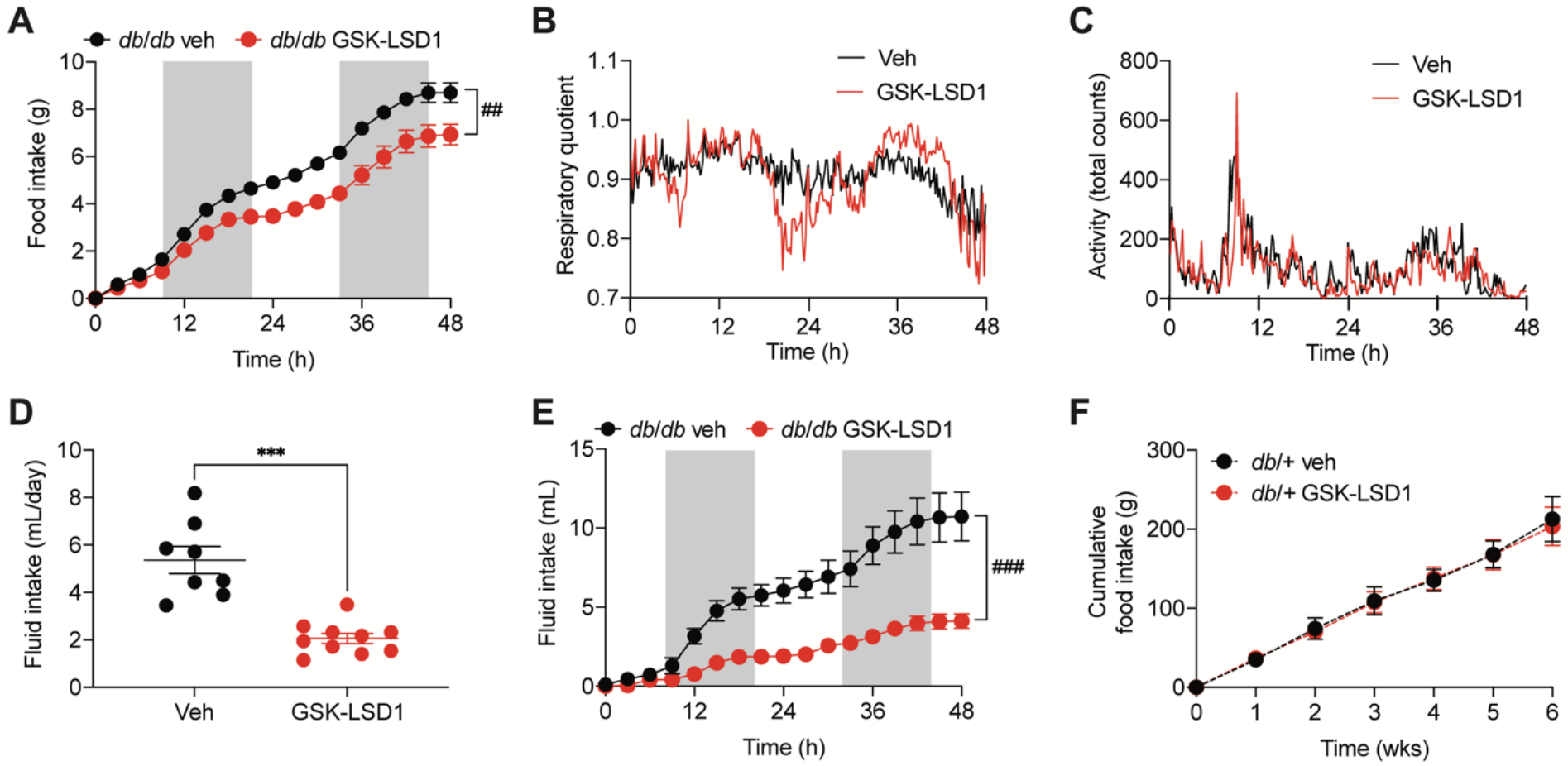
LSD1 inhibition does not alter energy expenditure in *db*/*db* mice. (**A**) Mice were injected daily with GSK-LSD1 or vehicle (veh) for 5 weeks and then placed into metabolic cages. Using the Comprehensive Laboratory Monitoring System (CLAMS), food intake was monitored and is shown as additive food intake over 48 hours (n = 4-5 mice/group). (**B**) Respiratory quotient over the course of 48 hours (n = 4-5 mice/group). (**C**) Spontaneous motor activity along the X-, Y-, and Z-axis with IR photocells (n = 4-5 mice/group). (**D**) Daily water consumption (n = 4-5 mice/group/day) and (**E**) Cumulative water consumption (n = 4-5 mice/group). (**F**) Cumulative food intake in lean *db*/+ mice treated with GSK-LSD1 or veh for 6 weeks (n = 3 cages/group, total n = 7-9 mice/group). Data presented as mean ± SEM. Statistical differences were calculated using an unpaired Student’s t-test (C) or two-way ANOVA (A, B, D-F) with Tukey post hoc analysis. \**p*<0.05, \*\**p*<0.01, \*\*\**p*<0.001.

**Figure S6.**
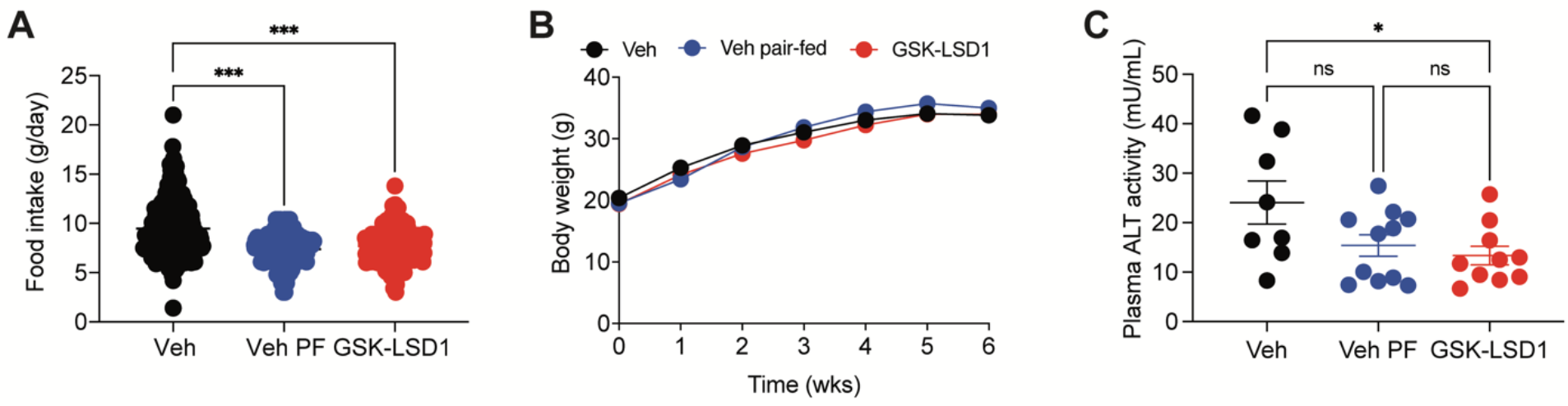
Metabolic parameters of pair-feeding study in *db*/*db* mice. (**A**) 4-week-old *db*/*db* mice were injected daily with vehicle (veh) or GSK-LSD1 for 6 weeks and fed a normal chow diet ad libitum. A third group of mice received veh and was pair-fed to GSK-LSD1-treated mice. Food intake was measured daily (n = 8-11 mice/group/day). (**B**) Body weight measured weekly (n = 8-11 mice/group). (**C**) Plasma ALT activity (n = 8-11 mice/group). Data presented as mean ± SEM. Statistical differences were calculated using a one-way ANOVA (A, D) or two-way ANOVA (B, C) with Tukey post hoc analysis. \**p*<0.05, \*\**p*<0.01, \*\*\**p*<0.001, ns, not significant.

**Figure S7.**
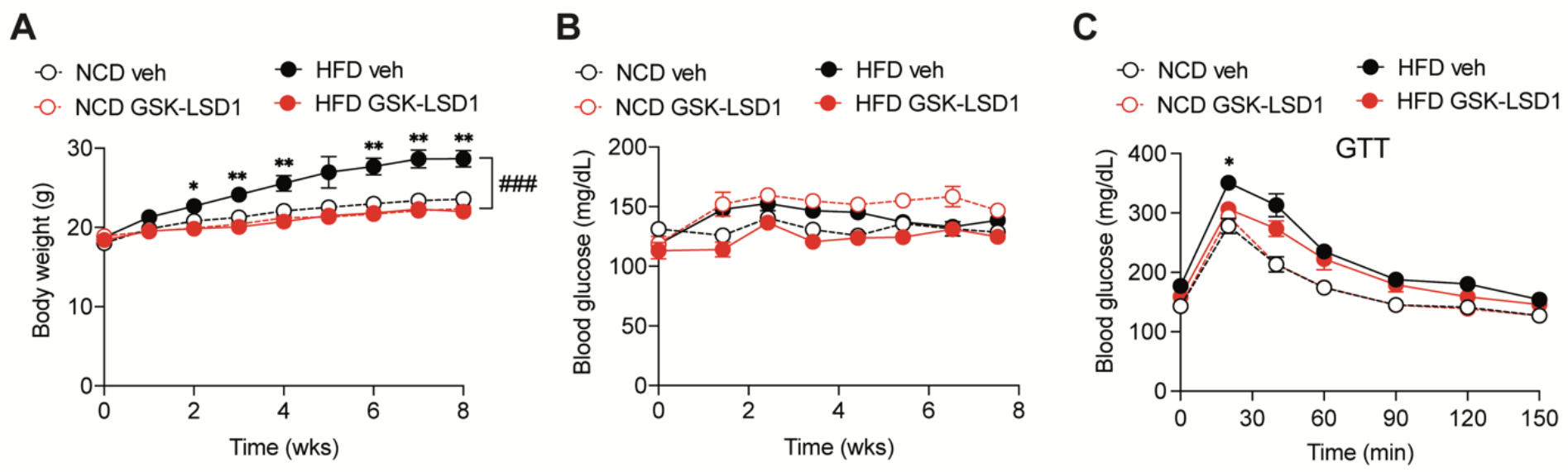
LSD1 inhibition prevents weight gain in high fat diet-fed mice. (**A**) 10-week-old C57BL/6J *wild type* (*WT*) mice were fed a high fat diet (HFD) or normal chow diet (NCD) for 8 weeks and injected daily with GSK-LSD1 or vehicle (veh). (**B**) Body weight and (**C**) blood glucose levels measured weekly (n = 8 mice/group). Asterisks indicate statistical differences between HFD GSK-LSD1- and HFD veh-treated mice. (**C**) Blood glucose levels after 7 weeks of treatment at indicated time points after a glucose bolus via oral gavage (n = 8 mice/group). Glucose tolerance test, GTT. Asterisk indicates statistical difference between HFD GSK-LSD1 and HFD veh mice. Data presented as mean ± SEM. A two-way ANOVA with Tukey post hoc analysis was performed to determine statistical differences. \**p*<0.05, \*\**p*<0.01, ^###^*p*<0.001.

**Figure S8.**
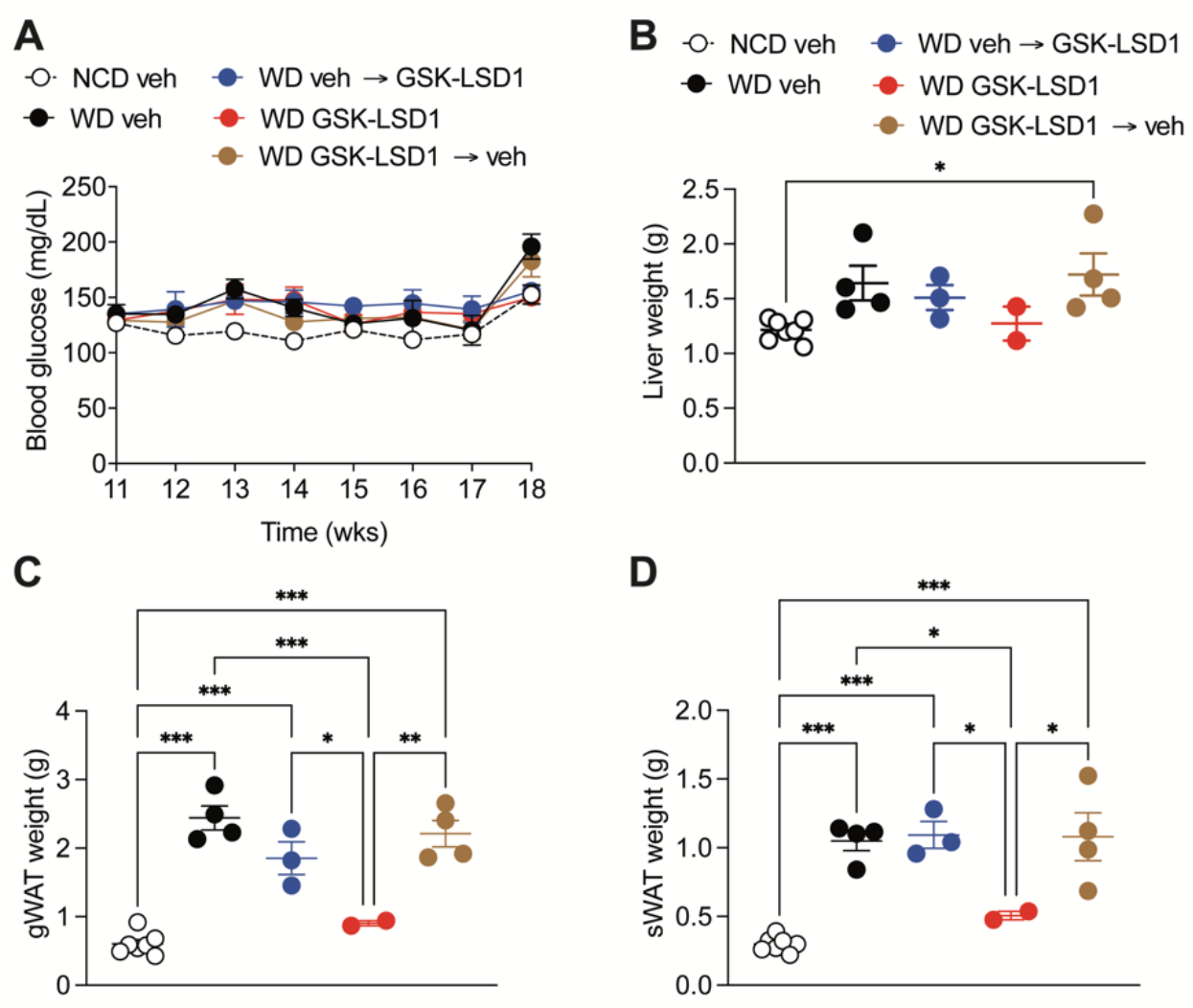
The effect of GSK-LSD1 on body weight in Western diet-fed mice is reversible. (**A**) 10-week-old C57BL/6J mice fed a Western diet (WD) were injected daily with GSK-LSD1 or vehicle (veh) for 11 weeks. As a control, a group of mice received veh and was kept on a normal chow diet (NCD). After 11 weeks of veh administration, the veh group was split into one group continuing veh administration (black), whereas the other half began to receive GSK-LSD1 daily for another 7 weeks (blue). Likewise, the GSK-LSD1 group was split into one group continuing GSK-LSD1 administration after 11 weeks (red), whereas the other half began to receive veh daily for another 7 weeks (brown). Blood glucose levels were measured weekly (NCD veh: n = 7 mice, WD groups: n = 4 mice). (**B-D**) Tissue weights for (**B**) liver, (**C**) gWAT, and (**D**) sWAT (n = 2-7 mice/group). Data are shown as mean ± SEM. Statistical differences were calculated using a two-way ANOVA (A-D) with Tukey post hoc analysis. \**p*<0.05, \*\**p*<0.01, \*\*\**p*<0.001.

**Supplemental Table S1.**
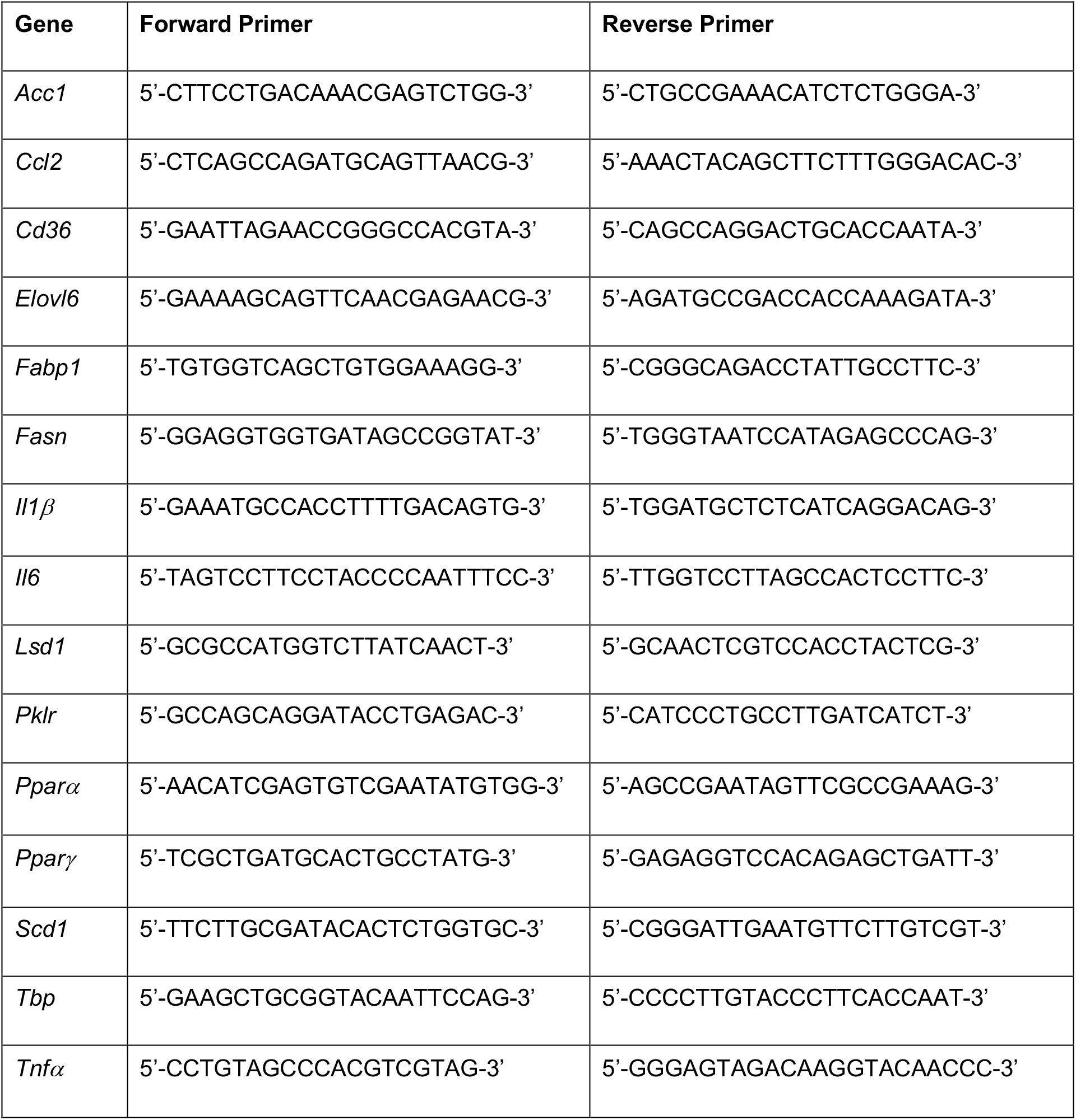
List of quantitative PCR primers.

